# Delta/Notch signaling controls neuroepithelial morphogenesis in the zebrafish spinal cord

**DOI:** 10.1101/517714

**Authors:** Priyanka Sharma, Vishnu Muraleedharan Saraswathy, Li Xiang, Maximilian Fürthauer

## Abstract

The morphogenesis of the nervous system requires coordinating the specification and differentiation of neural precursor cells, the establishment of neuroepithelial tissue architecture and the execution of specific cellular movements. How these aspects of neural development are linked is incompletely understood. Here we inactivate a major regulator of embryonic neurogenesis - the Delta/Notch pathway - and analyze the effect on zebrafish central nervous system morphogenesis. While some parts of the nervous system can establish neuroepithelial tissue architecture independently of Notch, Notch signaling is essential for spinal cord morphogenesis. In this tissue, Notch signaling is required to repress neuronal differentiation and promote neuroepithelial apico-basal polarity. Concomitant with a loss of their neuroepithelial properties, Notch signaling deficient cells also alter their morphogenetic behavior. In the wild-type zebrafish neural tube, cells divide at the organ midline to contribute one daughter cell to each organ half. Notch deficient animals fail to display this behavior and therefore form a misproportioned spinal cord. Taken together, our findings show that Notch signaling governs not only the cellular composition but also the morphogenetic shaping of the zebrafish spinal cord.

## INTRODUCTION

The building of functional organs requires controlling the identity, the shape and the spatial arrangement of their constituent cells. Understanding how these aspects of embryogenesis are linked remains a major challenge. The Delta/Notch pathway governs the specification, proliferation, and differentiation of neuronal precursors in both embryonic and adult tissues^1–5^. Notch receptors and Delta ligands are transmembrane proteins that elicit signaling between adjacent cells. In this context, the E3-Ubiquitin ligases Mindbomb and Neuralized promote an endocytic internalization of Delta ligand molecules that is required for Notch receptor activation^6–8^. Delta/Notch interactions then trigger a metalloprotease-mediated cleavage in the Notch extracellular domain, followed by a γ-Secretase-dependent intramembrane proteolysis that releases the Notch IntraCellular Domain (NICD) into the cytoplasm. NICD enters the nucleus to interact with CSL (for CBF1, Suppressor of Hairless, Lag1) family transcription factors and promote target gene transcription^1,4^.

Several observations suggest that Notch signaling and neuroepithelial morphogenesis are functionally interdependent. Notch has notably been linked to the formation of radial glia^9–12^, neural progenitor cells that present hallmarks of apico-basal polarity^13^. At the onset of neurogenesis, the neural plate of tetrapod embryos consists of a pseudostratified monolayer of apico-basally polarized cells which act as neural stem cells^13,14^. Following an expansion of this stem cell pool through symmetric divisions, some neural plate cells divide asymmetrically to generate the first neurons. Concomitant with this onset of neurogenesis, neural plate cells transform into radial glia^13^. During further development, radial glia cells undergo either symmetric, self-renewing divisions or divide asymmetrically to ultimately generate the majority of neurons that are present in the nervous system^13^. As radial glia cells retain most features of epithelial polarity, their presence is essential to maintain the epithelial architecture of the developing neural tube^10,13,15^.

In the forebrain of mice and zebrafish, cells undergoing neuronal differentiation present Delta ligands to activate Notch and maintain radial glia identity in neighboring progenitors^10,11^. Conversely, the apico-basal organization of the developing neural tube is itself required for Notch signaling as apical adherens junctions between nascent neurons and undifferentiating progenitors are required for receptor activation^15,16^.

In contrast to tetrapods, the initial stages of zebrafish spinal cord morphogenesis take place in a neural primordium that lacks a polarized epithelial architecture^17–21^. While the cellular organization of the zebrafish neural plate displays similarities to the pseudostratified epithelium of higher vertebrates^14,18,21,22^, major signs of apico-basal polarity such as apical Par protein localization, adherens junctions and tight junctions appear only by mid-segmentation stages, when neurogenesis is well under way^17-21,23^. This raises the question whether and how Notch signaling and neuroepithelial morphogenesis are linked during zebrafish neural development?

A second particularity of the development of the zebrafish is the occurrence of a particular type of morphogenetic cell division^17,18,20,24^. In C-divisions, a cell originating from one side of the neural primordium divides at the embryonic midline so that one of its daughters integrates the contralateral half^17,20,24^. The apical polarity protein Partitioning defective 3 (Pard3) accumulates at the cytokinetic bridge which prefigures the future apical neural tube midline^20^. Neural primordia in which cell divisions have been blocked establish apico-basal polarity, but fail to form a straight, regular tube apical neural tube midline^25^. It has therefore been suggested that C-divisions, while not being absolutely required for the establishment of neural tube apico-basal polarity, confer a morphogenetic advantage to the embryo by relocating cells that would otherwise span the neural tube midline^18,21,25^. Despite the fact that C-divisions confer robustness to neural tube development, their regulation remains poorly understood. While Pard3 and Planar Cell Polarity (PCP) proteins are known to control C-divisions^20,24^, their relationship to neurogenic Notch signaling has not been investigated.

Here, we inhibit Notch pathway activity and study the impact on zebrafish Central Nervous System (CNS) morphogenesis. Our work reveals that the relationship between Notch signaling and neuroepithelial morphogenesis depends on the biological context. While some regions of the nervous system can acquire apico-basal polarity and neuroepithelial organization independently of Notch, Notch signaling is required for the morphogenesis of the dorso-medial spinal cord. In this tissue, Notch signaling is essential to inhibit neuronal differentiation and promote the specification of neuroepithelial identity to allow the progressive epithelialization of the developing neural tube. Loss of Notch signaling also impairs the morphogenetic behavior of the cells of the neural primordium, thereby causing the formation of a misproportioned spinal cord. Our findings therefore show that, in addition to controlling neurogenesis, the Delta/Notch pathway plays a major role in the morphogenetic shaping of the zebrafish spinal cord.

## RESULTS

### Mindbomb1 is essential for zebrafish spinal cord morphogenesis

Genetic studies in the zebrafish have identified the E3-ubiquitin ligase Mindbomb1 (Mib1) as a central regulator of Delta ligand internalization and Notch activation^6,26^. To study the role of Notch signaling in zebrafish CNS morphogenesis, we inactivated *mib1* using a validated Morpholino^27^ and *mib1^ta52b^* mutants^6^. In accordance with published observations^27^, mib1 morphants presented an upregulation of DeltaD (DlD) due to a failure in Notch-dependent lateral inhibition, and a relocalization of DlD from endocytic compartments to the plasma membrane (Fig. 1a”,b”).

**Figure 1:**
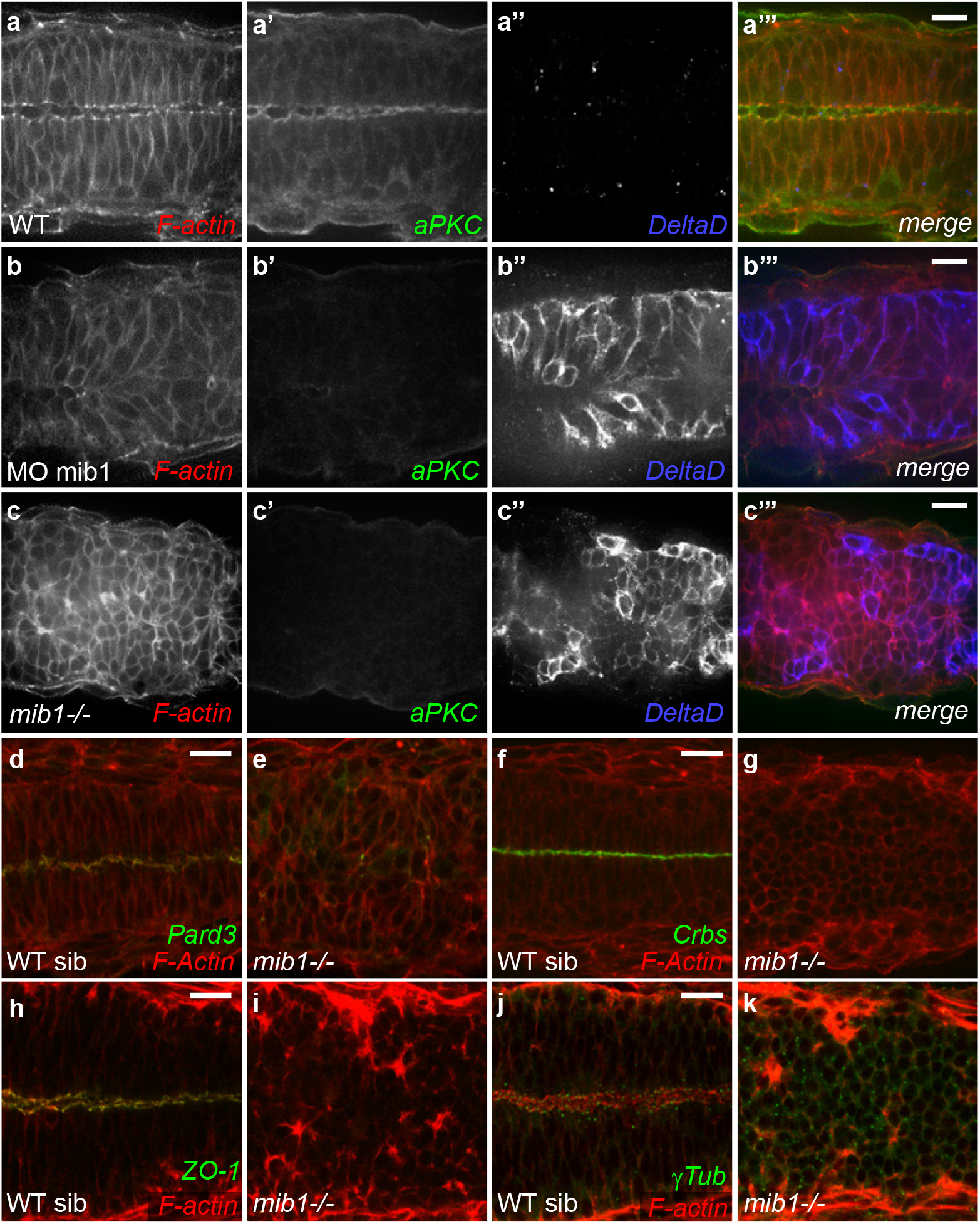
Mindbomb1 is required for zebrafish spinal cord morphogenesis. (a,b) Morpholino knock-down of mib1 disrupts apical aPKC enrichment and neuro-epithelial morphology (outlined by F-actin staining) in 15/19 embryos. (c) A similar disruption of apico-basal polarity is observed in *mib1* mutants (n=14). (d,e) *mib1* mutants fail to establish polarized Pard3 localization (e, n=12). (f,g) No polarized Crumbs enrichment is observed in *mib1* mutants (g, n=17). (h,i) The apical localization of the tight junction component ZO-1 is disrupted in *mib1* mutants (i, n=16). (j,k) Centrosomes fail to move towards the neural tube midline in *mib1* mutants (k, n=16). a-c,f,g, 30 somites stage. d,e, 18 somites stage, h-k 22 somites stage. All images are dorsal views of the anterior spinal cord, anterior left. Scalebars: 20 μm.

Antibody staining against the apical Par complex component atypical Protein Kinase C (aPKC^28^) was used to visualize apico-basal polarity. To analyze tissue morphology, we used fluorescent Phalloidin to visualize cortical F-actin. mib1 morphants display a loss of apical aPKC signal and an overall disorganization of neuro-epithelial tissue architecture in the anterior spinal cord (Fig. 1b,b’). A similar loss of neuroepithelial polarity was observed in *mib1^ta52b^* mutants (Fig. 1c,c’). Wild-type mib1 RNA injection rescued the polarity defects of *mib1^ta52b^* mutants, warranting the specificity of our observations (Supplementary Fig. S1).

Our experiments show that loss of Mib1 impairs apico-basal polarity and epithelial organization in the zebrafish spinal cord. Accordingly, no polarized enrichment of the apical polarity proteins Pard3^29^ (Fig. 1d,e), Crumbs^30^ (Fig. 1f,g) and the junctional component Zonula Occludens 1 (ZO-1^17^, Fig. 1h,i) is detectable in *mib1^ta52b^* mutants. In accordance with a failure to establish Par complex-dependent polarity, *mib1^ta52b^* mutants fail to display the Pard3-dependent alignment of γTubulin-positive centrosomes that is observed at the neural tube midline of wild-type siblings (Fig. 1j,k)^31^.

### Canonical Notch signaling is required for the morphogenesis of the zebrafish spinal cord

In addition to its role in Delta ligand internalization, Mib1 also regulates the ubiquitination and endocytic trafficking of other substrate proteins^32,33^. This raises the question whether the neuroepithelial defects of Mib1-depleted embryos are due to the loss of Notch signaling or to a Notch-independent function of Mib1? To address this issue, we interfered with different Notch pathway components and analyzed the effect on spinal cord morphogenesis.

Mib1 loss-of-function impairs the endocytosis of DlD, one of the two zebrafish homologues of mammalian Delta-like-1 (Dll1) ^6,27^. Apico-basal polarity is however intact in *dld^ar33^* mutants (Supplementary Fig. S2a,b). Mib1 also interacts with DeltaA (DlA), the second Dll1 homologue^34^. Accordingly, *mib1^ta52b^* mutants display excessive cell surface accumulation of DlA (Fig. 2a,b). Injection of a validated dla morpholino^35^ abolishes DlA immunoreactivity but fails to elicit polarity defects in a wild-type background (Supplementary Fig. S2c,d). In contrast, polarity defects are observed upon dla knock-down in *dld^ar33^* mutants (Fig. 2c,d).

**Figure 2:**
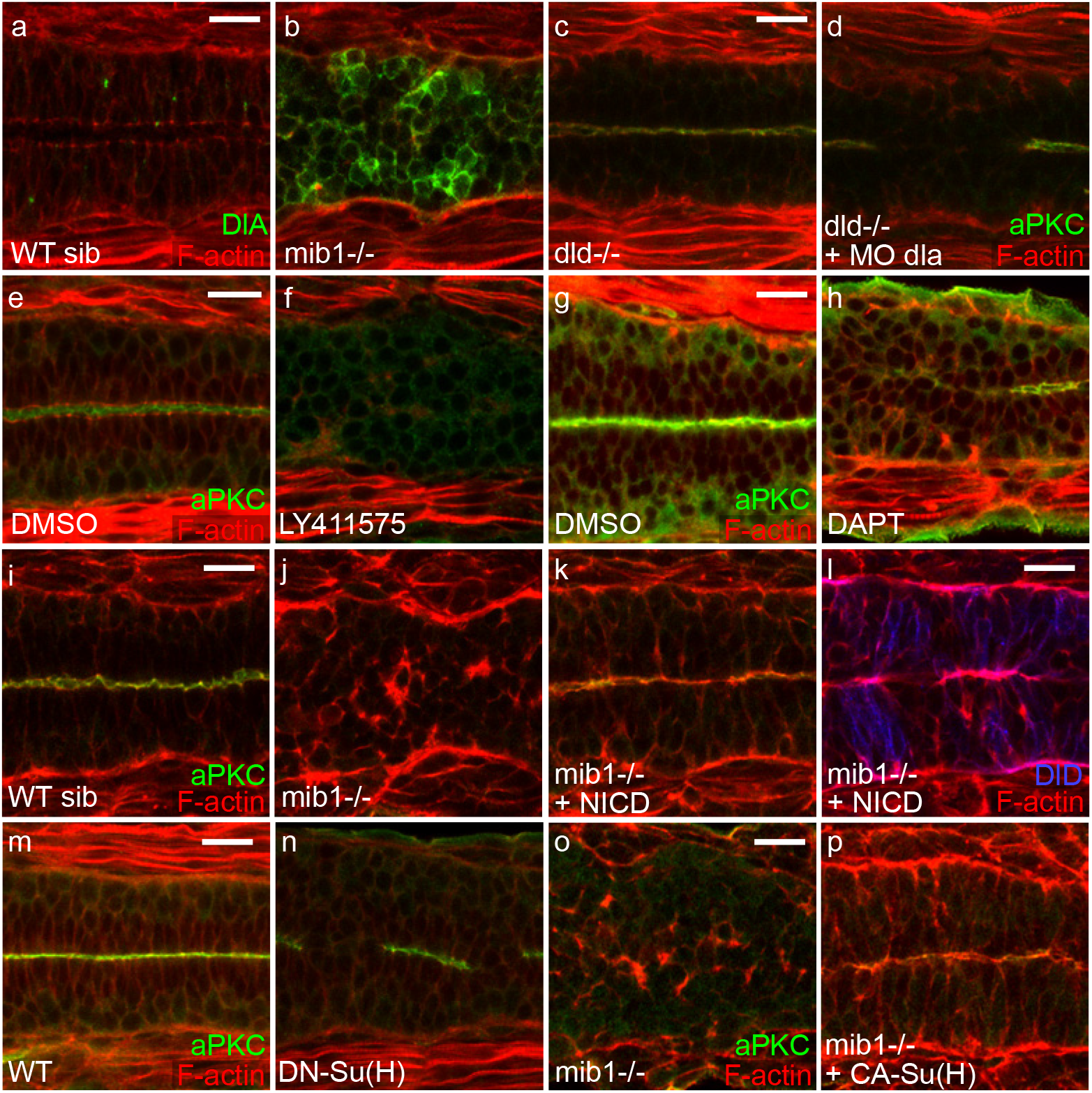
Notch pathway activity is required for spinal cord morphogenesis. (a,b) *mib1* loss of function prevents DlA internalization (n=9). (c,d) Combined inactivation of *dld* and dla disrupts apico-basal polarity in 9/10 embryos. (e,f) The γ-Secretase inhibitor LY411575 disrupts apico-basal polarity (n=18). (g,h) Similarly the γ-Secretase inhibitor DAPT perturbs polarity in 4/5 embryos. (i-l) RNA injection of a constitutively activated form of Notch (NICD) restores neuro-epithelial morphology and apical aPKC localisation (k) but not DeltaD endocytosis (l) in 22/22 embryos. (m,n) Polarity defects are observed in 20/46 embryos injected with RNA encoding dominant-negative Su(H) (DN-Su(H)). (o,p) RNA injection of constitutively activated Su(H) (CA-Su(H)) restores apico-basal polarity in 17/17 *mib1* mutants. All images are dorsal views of the anterior spinal cord, anterior left. a-f 22 somites stage, g,h,m,n 30 somites stage, i-l,o,p 16 somites stage. Scalebars: 20 μm.

Notch receptor activation results in the γ-Secretase-mediated release of NICD into the cytoplasm, allowing NICD nuclear entry and transcriptional activation of target genes^1,4^. Blocking Notch signaling using two different pharmacological γ-Secretase inhibitors, DAPT^36^ and LY411575^37^, impaired the apico-basal polarization of the neural tube (Fig. 2e-h).

If the polarity phenotype of Mib1 depleted embryos is a consequence of the loss of Notch signaling, restoring Notch activity should rescue neuroepithelial morphogenesis. Accordingly RNA injection of NICD, which acts as a constitutively activated form of Notch^38^, restores neural tube apico-basal polarity in *mib1^ta52b^* mutant (Fig. 2i-l) or mib1 morphant (Supplementary Fig. 2e-h) embryos.

Upon nuclear entry NICD associates with RBPJ/Su(H)/CBF transcription factors to trigger target gene activation^1,4^. Misexpression of a dominant-negative Su(H) variant^39^ impaired neuroepithelial morphogenesis and polarized aPKC localization (Fig. 2m,n), consistent with the phenotype of CBF1 knockout mice^40^. This result was confirmed through simultaneous morpholino knock-down of the two zebrafish Su(H)-homologues RBPJa&b (Supplementary Fig. 2i-l)^41^. Conversely, RNA microinjection of Constitutively Activated Su(H) (CA-Su(H)^39^) restores neuroepithelial tissue organization in *mib1^ta52b^* mutants (Fig. 2o,p).

Our observations therefore show that not only Mib1 itself, but the activity of the entire canonical Notch pathway is required for zebrafish spinal cord morphogenesis.

### Notch signaling is dispensable for the early establishment of floor plate apico-basal polarity

Previous studies have suggested that canonical Notch signaling is dispensable for the initial establishment, but required for the subsequent maintenance of neuroepithelial apico-basal polarity in fish and mice^40,42^. To address whether the defects of Mib1-depleted embryos similarly arise from a failure to maintain neuroepithelial polarity, we monitored the establishment of neural tube apico-basal polarity in wild-type sibling and *mib1^ta52b^* mutant embryos.

The emergence of polarity has been studied mostly in the dorsal and medial regions of the zebrafish spinal cord^17,20^. In accordance with these studies, we find that the dorso-medial spinal cord does not show overt signs of apico-basal polarity prior to the 12 somites stage. We noticed however that apico-basal polarity emerges much earlier in the ventral-most part of neural tube. From the 6 somites stage onwards, embryos display foci of polarized aPKC expression (Fig. 3a,c) that coalesce subsequently into a continuous line (Fig. 3g’, Supplementary Fig. 3). Lateral views of the neural tube show that this enrichment of aPKC (Fig. 3c) and Crumbs (Fig. 3g, Supplementary Fig. 3) corresponds to the apical surface of the cuboidal cells of the floor plate.

**Figure 3:**
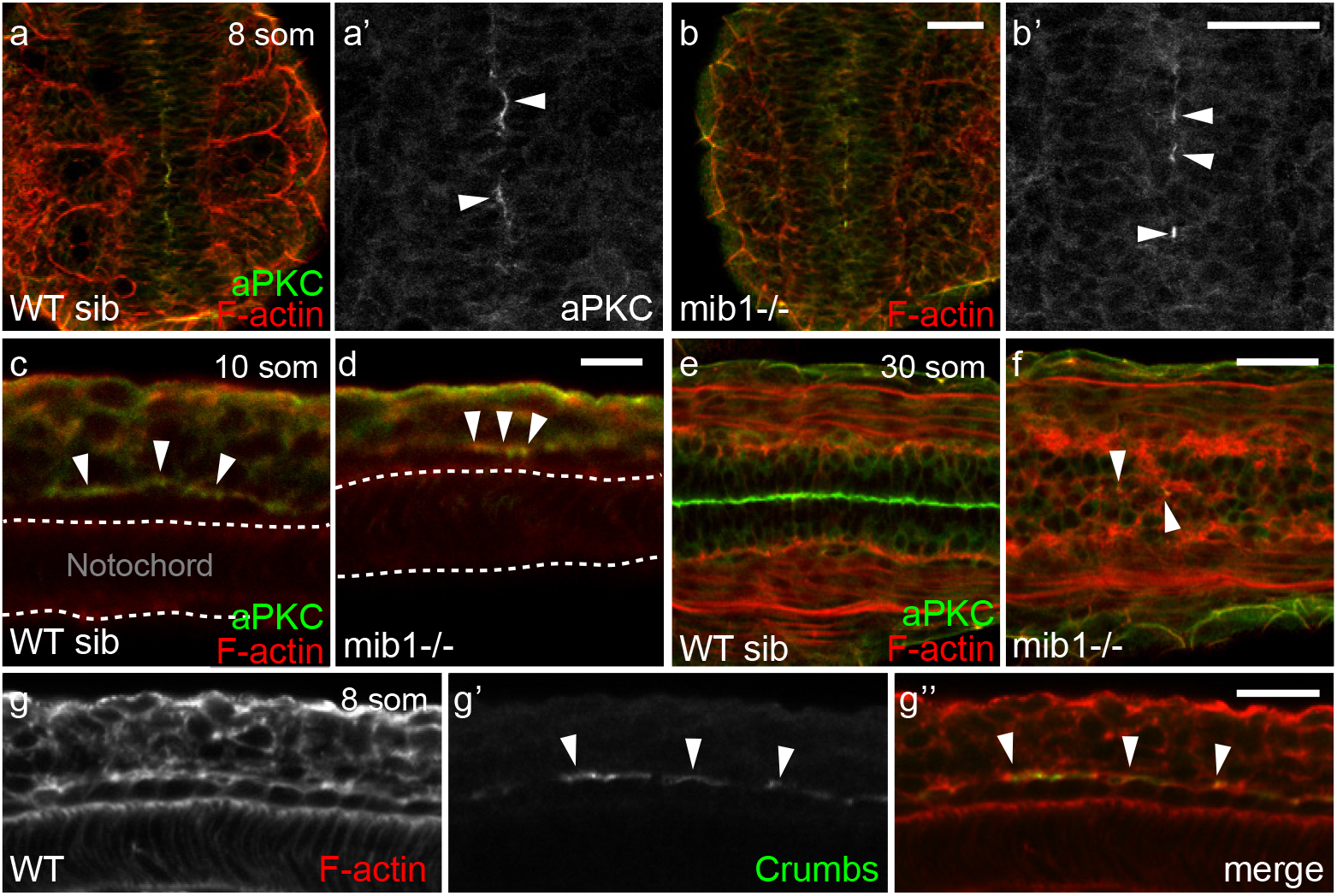
Notch signaling is dispensable for the emergence of floor plate apico-basal polarity. (a,b) At the 8 somites stage, similar discontinuous patches of polarized aPKC are detected in the ventral-most neural tube of WT siblings (a, n=31) and *mib1* mutants (b, n=14). Dorsal views, anterior up. (a’,b’) are high magnification views of the polarized aPKC signal. (c,d) aPKC is enriched at the apical surface of floor plate cells in 10 somites stage WT siblings (c, n=8) and *mib1* mutants (d, n=4). Lateral views, anterior left. (e,f) By the 30 somites stage polarized aPKC staining is reduced to few isolated cells in the ventral-most neural tube of *mib1* mutants (arrowheads in f, n=5). Dorsal view, anterior left. (g) Crumbs protein accumulates at the apical surface of 8 somites stage floor plate cells (arrowheads). Lateral view, anterior left. Scalebars: (a,b) 40μm, (c,d) 10 μm, (e-g) 20 μm.

Notch signaling has been implicated in the differentiation of floor plate cells^43^. Mutations in *dla* or *mib1* have been reported to cause a severe reduction in the number of detectable floor plate cells by the end of the segmentation period^43^. Our observations confirm the occurrence of late floor plate defects (Fig. 3e,f), but reveal that the initial establishment of floor plate cells and their apico-basal polarization do not require *mib1* function (Fig. 3a-d) and may therefore occur independently of Notch signaling. To test this hypothesis, we took advantage of the transgenic *tp1:bglob-GFP* Notch reporter line^44^. At the stages considered here, reporter activity was absent from the floor plate, while adjacent tissues displayed fluorescence indicative of Notch signaling (Supplementary Fig. 3). This observation confirms that Notch signaling is dispensable for the initial formation and apico-basal polarization of the ventral-most cells of the neural tube.

### Notch controls neuroepithelial tissue architecture in the dorso-medial spinal cord

In contrast to the situation in the floor plate, Notch is critically required for the morphogenesis of the more dorsal regions of the spinal cord (Fig. 4). To analyze the emergence of neuroepithelial tissue architecture and apico-basal polarity in the anterior spinal cord, we performed a time course analysis of aPKC localization. In wild-type controls, aPKC becomes progressively enriched at the neural tube midline in the medial and dorsal aspects of the spinal cord from the 12 somite stage onwards (Fig. 4b,f,j,m,n). In contrast, *mib1^ta52b^* mutants fail to display neuroepithelial tissue architecture and apico-basal polarity in the dorso-medial spinal cord at all stages examined (Fig. 4d,h,l,m,o). While previous studies have highlighted functions of Notch signaling in the late maintenance of neurectodermal apico-basal polarity^40,42^, our findings show that zebrafish *mib1* is required already to initiate the establishment of neuroepithelial tissue architecture.

**Figure 4:**
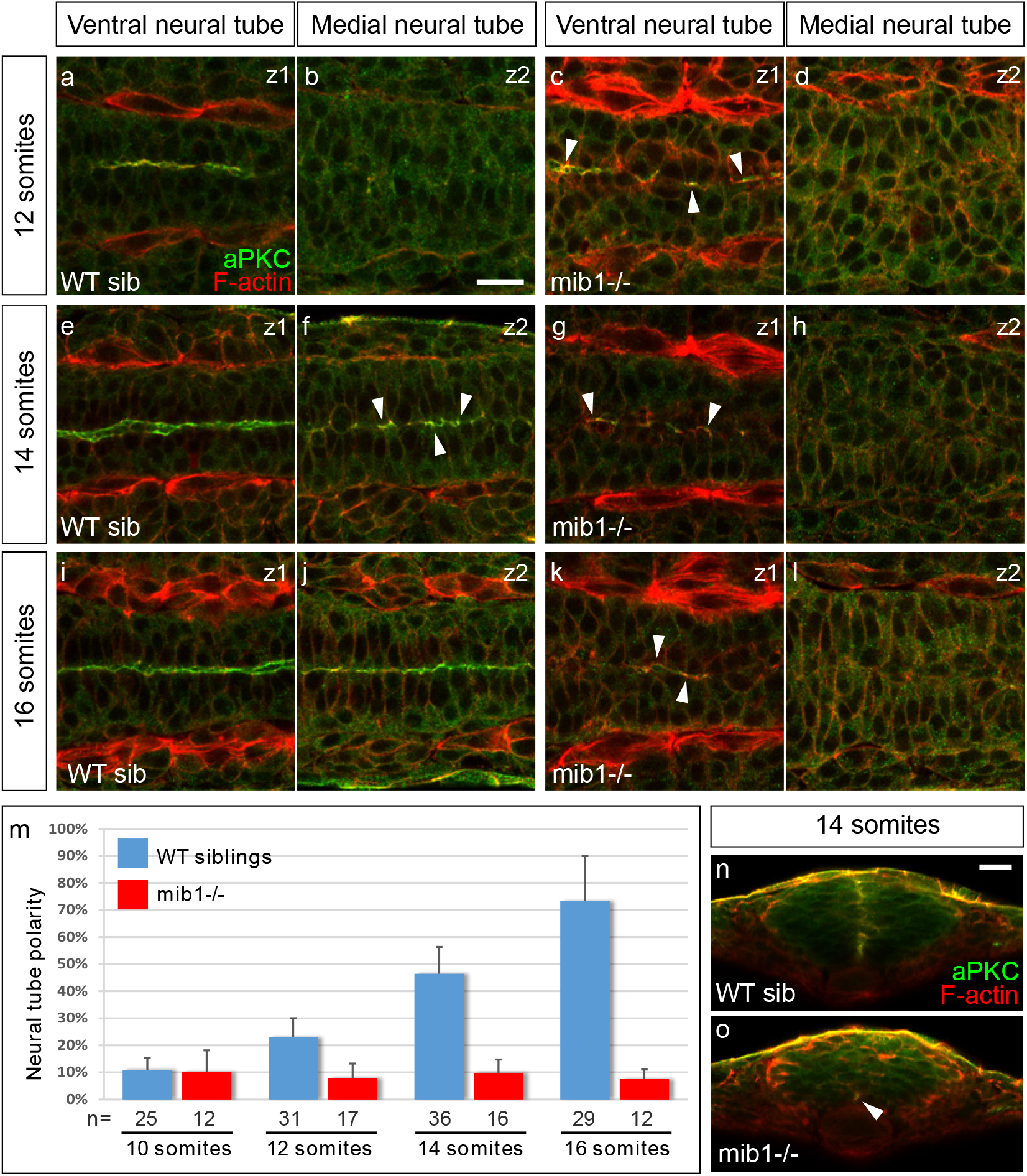
Notch signaling is required for the establishment of apico-basal polarity in the dorso-medial spinal cord. (a-l) Confocal sections taken at different dorso-ventral levels of the anterior spinal cord of WT sibling and *mib1* mutant embryos. Dorsal views, anterior left. z1 corresponds to the ventral-most extent of apico-basally polarized neuro-epithelial tissue (identified by aPKC staining), z2 is localized 12 μm more dorsally in the same embryo. Arrowheads indicate local foci of polarized aPKC in partially polarized tissue. (a-d) At the 12 somites stage polarized aPKC signal is detected in the ventral-most neural tube in WT sibling and *mib1* mutants. (e-l) At later stages polarity is progressively established in more dorsal regions of the neural tube in WT siblings (f,j), but remains limited to the ventral neural tube in *mib1* mutants (g,h,k,l). (m) Quantification of the progressive emergence of apico-basally polarity in the anterior spinal cord (see Methods). Error bars represent SD. (n,o) Transversal sections (dorsal up) through the neural tube of 14 somites stage embryos. (n) Polarized aPKC staining starts to spread through the dorso-ventral extent of the neural tube in WT siblings. (o) In *mib1* mutants polarized aPKC enrichment remains limited to the ventral floor plate region (arrowhead). Scalebars: 20 μm.

Towards the end of neural development, most neural progenitors downregulate the expression of apical polarity proteins and differentiate into neurons^45^. As inactivation of *mib1* causes premature neuronal differentiation^6^, we wondered whether the loss of neuroepithelial tissue organization in *mib1^ta52b^* mutants might be correlated with a failure to express genes governing neuroepithelial identity and apico-basal polarity. Zebrafish *sox19a* is expressed in undifferentiated neural precursors cells throughout the nervous system, similar to the expression of amniote sox2^46^. In accordance with a loss of neuronal precursors due to excessive neuronal differentiation, *mib1^ta52b^* mutants display a premature loss of *sox19a* expression in the spinal cord (Fig. 5d). Accordingly, all cells of the dorso-medial spinal cord start to express the marker of neuronal differentiation *elavl3* (Supplementary Fig. 4c’,c”).

**Figure 5:**
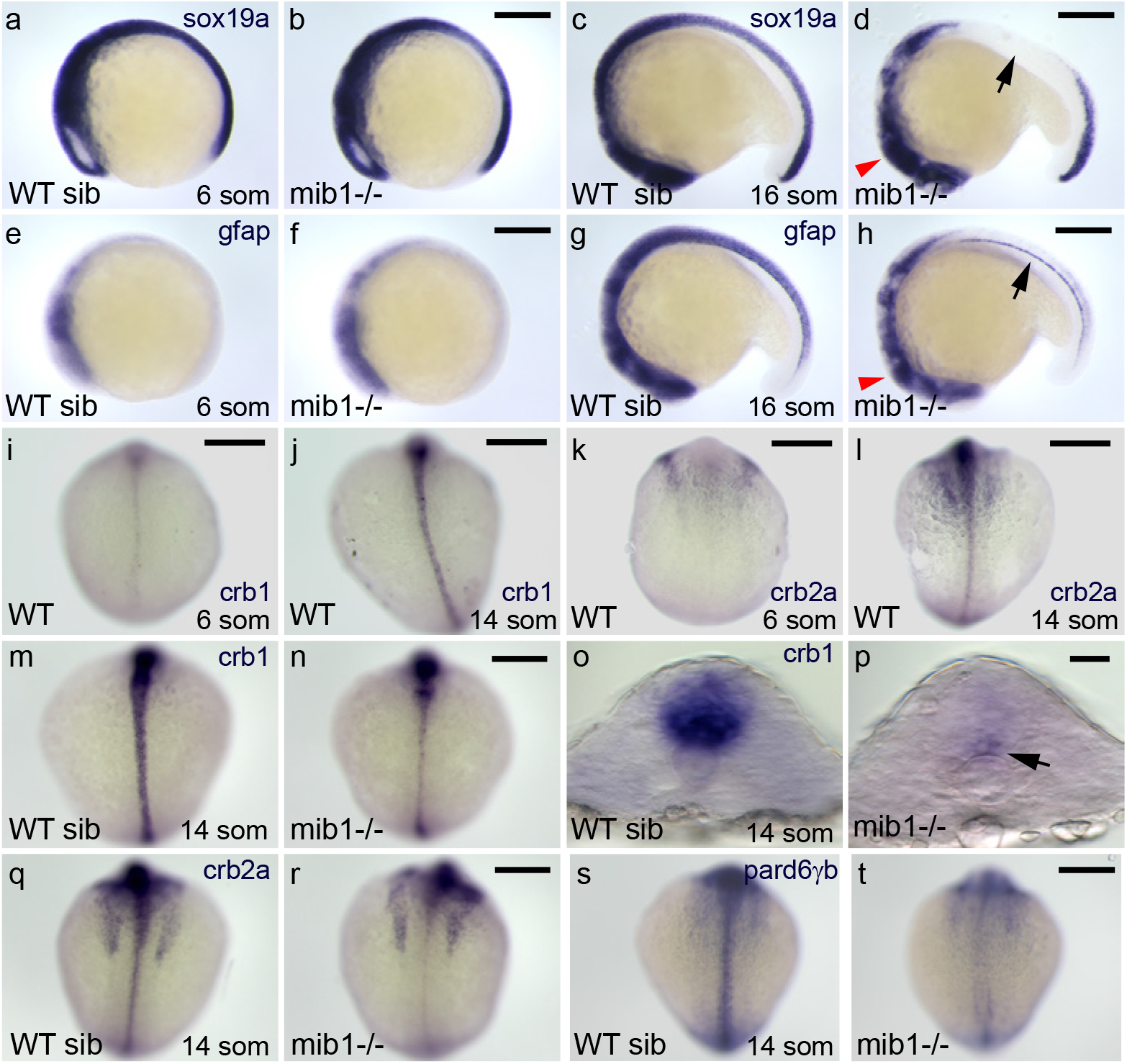
Notch signaling specifies neuroepithelial identity. (a,b) The expression of the neuronal progenitor marker *sox19a* is similarly initiated in WT sibling (a) and *mib1* mutant embryos (b, n=4). (c) By the 16 somites stage, *sox19a* expression is still present in the brain and anterior spinal cord of WT siblings. (d) In *mib1* mutants (n=17) *sox19a* is lost in the anterior spinal cord (arrow) but partially retained in the brain (red arrowhead). (e,f) At the 6 somites stage, low levels of the radial glia marker *gfap* are detected in the brain region of WT siblings (e) or *mib1* mutants (f, n=10). (g) By the 16 somites stage, WT siblings display *gfap* expression in the brain and the spinal cord. (h) *mib1*mutants (n=17) fail to upregulate *gfap* expression in the dorso-medial spinal cord and present expression only in the floor plate (arrow) and brain (red arrowhead). (i-l) The establishment of apico-basal polarity coincides with an upregulation of *crb1* (i,j) and *crb2a* (k,l) in the spinal cord. (m,n) Upregulation of *crb1*expression is impaired in *mib1* mutants (n, n=11). (o,p) Residual *crb1* expression persists in the ventral-most spinal cord (arrow in p) of 14 somites stage *mib1* mutants. (q,r) *mib1* mutants display reduced *crb2a*expression in the spinal cord (r, n=13). (s,t) *pard6γb* expression is lost in the neural tube of *mib1* mutants (t, n=13). a-h, lateral views, anterior to the left, dorsal up. i-n, q-t, dorsal views of the spinal cord, anterior up. o,p, transversal sections of the neural tube, dorsal up. Scalebars: (a-n,q-t) 200 μm, (o,p) 20 μm.

Radial glia cells are neural precursors that display apico-basal polarity and are crucial for the maintenance of neuroepithalial tissue architecture in the brain of zebrafish and higher vertebrates^10,11^. One of the hallmarks of radial glia is the expression of *glial fibrillary acidic protein (gfap)*^12,13^. In accordance with a lack of apico-basally polarized neural precursor cells, *gfap* expression levels are very low in the presumptive spinal cord of 6 somites stage embryos (Fig. 5e). While wild-type sibling embryos upregulate *gfap* expression concomitantly with the establishment of apico-basal polarity (Fig. 5g), *mib1^ta52b^* mutants fail to express *gfap* in the spinal cord (Fig. 5h).

Loss of Notch signaling impairs not only *gfap*, but also the expression of core components of the apico-basal polarity machinery belonging to the Par and Crumbs protein complexes^47^. While only low levels of *crumbs1 (crb1) and crumbs2a (crb2a)* are detectable in the neural tube of wild-type 6 somite stage embryos (Fig. 5i,k), both genes display increased expression by the 14 somites stage (Fig. 5j,l). *mib1^ta52b^* mutants fail to display this upregulation (Fig. 5m-r). Similarly, reduced expression levels of the Par complex component *pard6γb*^48^ are observed in the spinal cord of *mib1^ta52b^* mutants (Fig. 5s,t).

Already before the onset of neurogenesis, the neural plate of higher vertebrates displays hallmarks of epithelial organization. Our findings show that in zebrafish, the primordium of the developing spinal cord does initially not express markers of polarized neural precursor cells (*gfap*) and components of the apico-basal polarity machinery (*crb1, crb2a, pardbγB*). Lack of Notch signaling results in excessive neuronal differentiation and failure to upregulate the neuroepithelial gene expression program, explaining thereby why *mib1^ta52b^* mutants fail to display the progressive epithelialization that is observed in the neural tube of wild-type controls (Fig. 4m, Supplementary Fig. 4c).

### Region-specific control of neuroepithelial morphogenesis in the zebrafish spinal cord

In addition to uncovering an essential role of Notch in the regulation of neuroepithelial genes in the dorso-medial spinal cord, our analysis of gene expressions confirms the existence of a different, Notch-independent regulation of apico-basal polarity in the ventral-most part of the neural tube. In this tissue, *crb1* transcripts are detectable already by the 6 somites stage (Fig. 5i), when first signs of polarized aPKC enrichment become detectable. While *mib1^ta52b^* mutants fail to upregulate *crb1* expression in the dorso-medial neural tube where polarity is lost, *crb1* expression persists in the ventral-most cells where apico-basal polarity is retained (Fig. 5p). Similarly, *mib1^ta52b^* mutant floor plate cells retain the expression of the neuroepithelial/radial glia marker *gfap* (Fig. 5h, arrow). Accordingly, the analysis of the neuronal differentiation marker *elavl3* reveals that, in contrast to more dorsal and lateral spinal cord derivatives, *mib1^ta52b^* mutant floor plate cells do not undergo neuronal differentiation (Supplementary Fig. 4d)

In contrast to the spinal cord, *mib1^ta52b^* mutant brains do still express *gfap* and the neuronal precursor marker *sox19a*, albeit at reduced levels (arrowheads in Fig. 5d,h). Similarly, *crb1* and *crb2a* are still expressed in the brain but no more detectable in the dorso-medial spinal cord of 30 somites stage *mib1^ta52b^* mutants (Fig. 6a-d). The occurrence of *mib1^ta52b^* mutant polarity phenotypes correlates with the differential regulation of neuroepithelial gene expression. While apico-basal polarity is disrupted in the spinal cord of *mib1^ta52b^* mutants (Fig. 6h,l, Supplementary Fig. 5d), polarity is only partially disrupted in the hindbrain (Fig. 6j) and essentially unaffected at the Midbrain-Hindbrain Boundary (MHB) (Fig. 6b, Supplementary Fig. 5b). To determine whether the intact neuroepithelial organization of mutant MHBs could be due to residual Notch signaling, we introduced the *tp1bGlob:GFP* reporter^44^ into *mib1* mutants (Supplementary Fig.5). Notch reporter activity is essentially undetectable at the MHB in both wild-type sibling and homozygous mutant animals (Supplementary Fig. 5a,b). These observations suggest that Notch signaling is indeed largely dispensable for MHB development at the stages considered here.

**Figure 6:**
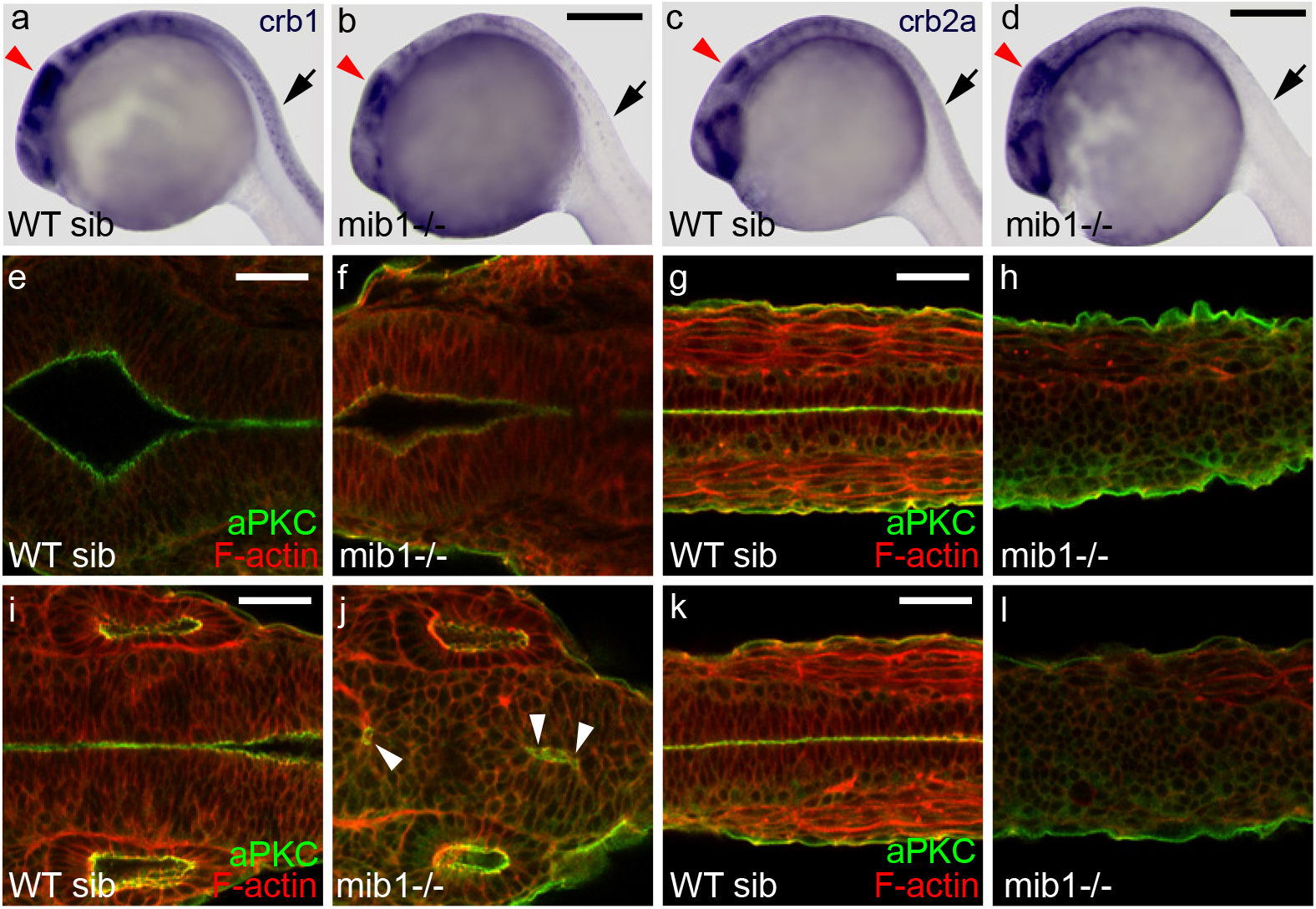
Notch differentially affects neuroepithelial polarity in the brain and spinal cord. (a-d) Lateral views (anterior left) of the head and anterior spinal cord at 30 somites show that in *mib1* mutants *crb1* (b, n=10) and *crb2a* (d, n=10) expression are lost in the anterior spinal cord (black arrows) but partially retained in the brain (red arrowheads indicate the midbrain). (e-l) In *mib1* mutants neuroepithelial apico-basal polarity is maintained at the level of the midbrain-hindbrain boundary (f, n=7), partially disrupted at the level of the hindbrain (j, arrowheads indicate residual polarized aPKC signal, n=7) but completely lost throughout the spinal cord (h,l). (k,l) represent the most anterior and (g,h) the trunk spinal cord. The picture pairs (e,g), (f,h), (i,k) and (j,l) each represent the same embryo imaged at different antero-posterior locations. e-l are dorsal views, anterior left. Scalebars: (a-d) 250 μm, (e-l) 40 μm.

### Apico-basal polarity does not require Notch signaling between midline-crossing mitotic sister cells

A characteristic feature of the morphogenesis of the zebrafish neural tube is the occurrence of midline-crossing C-divisions. As Notch signaling between mitotic sister cells is important for cell fate assignment during later stages of zebrafish neurogenesis^11,49^, we wondered whether Delta/Notch signaling between C-dividing sister cells might be important for apico-basal polarity?

To explore this possibility, *mib1^ta52b^* mutant cells were transplanted into wild-type hosts. As Mib1 is essential for Delta ligand activity, no Delta/Notch signaling occurs between *mib1^ta52b^* mutant mitotic sister cells. Despite this fact, mutant cells display a polarized localization of Pard3-GFP at the apical cell surface (Fig. 7a). Conversely, wild-type cells implanted into the dorso-medial spinal cord of *mib1^ta52b^* mutants hosts fail to display polarized aPKC localization (Fig. 7b). Spinal cord apico-basal polarity does therefore not require Delta/Notch signaling between midline-crossing mitotic sister cells.

**Figure 7:**
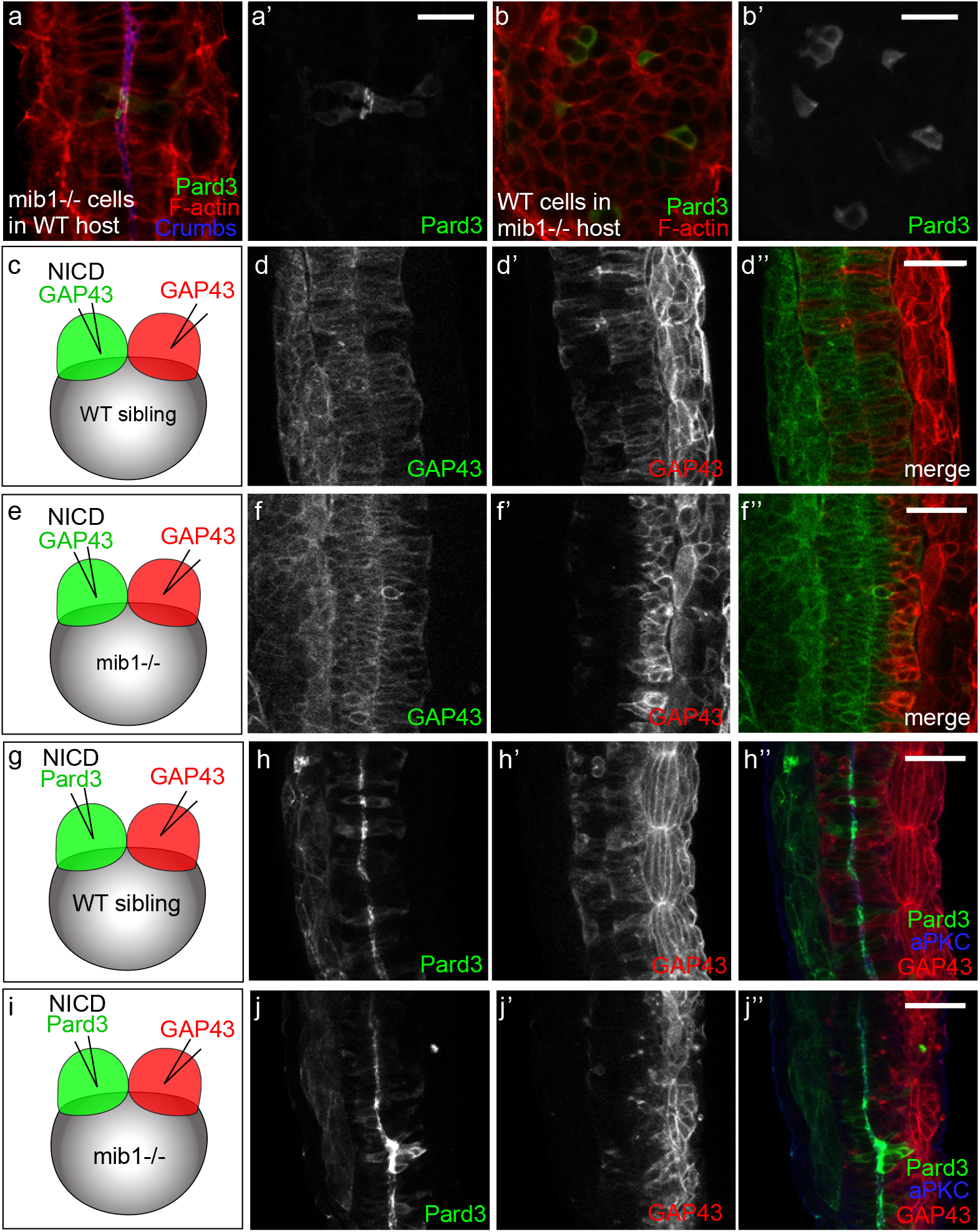
Dissection of the spatial requirement for Notch signaling in neural tube morphogenesis. (a,a’) Pard3-GFP expressing *mib1* mutant cells undergo correct polarization when transplanted into WT hosts (n=30 cells in 6 embryos). (b,b’) In contrast, Pard3-GFP expressing WT cells fail to polarize when transplanted into *mib1* mutant hosts (n=96 cells in 13 embryos). (a,b) are single confocal sections, (a’,b’) maximum projections of GFP-positive clones. (c-f) The two halves of the neural tube were labelled by injecting RNAs encoding red or green fluorescent membrane labels (GAP43) into the 2 blastomeres of 2cell stage embryos (see Methods). (c,d) In WT sibling embryos half-injected with RNA encoding constitutively activated Notch (NICD), cells originating from both sides of the neural tube cross the neural tube midline to integrate the contra-lateral organ half (n=7/7). (e,f) If NICD is half-injected into *mib1* mutants, NICD-containing cells display extensive midline crossing (f,f”), while the crossing of NICD-negative cells is reduced in 6/8 embryos (f’,f”). (g-j) One half of the embryo was injected with RNA encoding NICD Pard3-GFP, the other half with GAP43-RFP. (g,h) 6/6 WT sibling embryos display apical Pard3 accumulation (h,h”) and bilateral midline crossing (h’,h”). (i,j) In *mib1* mutants, NICD causes apico-basal polarization and midline crossing of Pard3-GFP positive cells in 6/6 embryos (j,j”). NICD-negative GAP43-RFP positive cells fail however to cross the neural tube midline in 5/6 embryos (j’,j”). Pictures represent dorsal views of the spinal cord (anterior up) at 30 somites (a,b), 18 somites (d,f) and 21 somites (h,j) stages. Scalebars: a,b 20 μm, d,f,h,j 40 μm.

### Cell autonomous Notch signaling is required for neuroepithelial morphogenesis

In the mouse and zebrafish forebrain, cells undergoing neuronal differentiation present Delta ligands to activate Notch in neighbouring cells and thereby maintain their radial glia identity^10,11,16^. If a similar mechanism is at work in the zebrafish spinal cord, Notch activity should govern the cell autonomous acquisition of neuroepithelial characteristics. To address this issue, we generated mosaic embryos in which Notch signaling is activated only in one half of the neural primordium (Fig. 7c-j, see Methods for details).

In a first set of experiments, RNAs encoding NICD and a fluorescent membrane label (GAP43-RFP) were co-injected into one blastomere of two cell stage embryos. A second injection was performed to introduce GAP43-GFP in the other blastomere (Fig. 7c,e). By the end of gastrulation embryos in which the progeny of the two injected blastomeres had populated the left and right sides of the animal were selected and grown further to analyze the morphology and behavior of cells in the neural primordium. In wild-type siblings, NICD-positive and NICD-negative cells originating from the two neural tube halves both adopted an elongated morphology with cell bodies spanning the apico-basal extent of the neuroepithelium (Fig. 7d,d’).

A different result was observed when the same manipulation was carried in *mib1^ta52b^* mutants (Fig. 7e,f). In this case, only NICD-positive cells adopted a characteristic epithelial morphology, while NICD-negative cells failed to contact the neural tube midline and populated the basolateral aspect of the neural tube, a behavior characteristic of neuronal differentiation. To confirm that NICD is able to promote the acquisition of apico-basal polarity, we performed a second set of experiments where NICD was co-injected with Pard3-GFP (Fig. 7g-j). These experiments confirmed that NICD is capable of promoting the apico-basal polarization of Pard3 in *mib^ta52b^* mutant cells (Fig. 7j).

Our observations suggest that Notch signaling is required cell autonomously for the acquisition of neuroepithelial characteristics. These experiments do however also provide evidence that, even in conditions where Notch signaling is active only in one half of the cells of the neural primordium (Fig. 7e,i), *mib1^ta52b^* mutant neural tubes present a continuous apical neural tube midline, as indicated by cell morphology (Fig. 7f), Pard3-GFP accumulation (Fig. 7j) and aPKC localization (Fig. 7j”). While Notch signaling promotes neuroepithelial characteristics only in the cells where it is active, the presence of a fraction of Notch-activating cells is therefore sufficient to convey overall neuroepithelial organization to the spinal cord.

### Notch signaling governs morphogenetic cell movements in the zebrafish spinal cord

In wild type zebrafish, midline-crossing C-divisions cause the intermingling of cells originating from the two halves of the neural tube (Fig. 7d)^17,20^. In experiments where Notch signaling was restored in one half of the *mib1^ta52b^* mutant neural primordium, we observed that only NICD-positive but not NICD-negative cells are able to cross the neural tube midline (Fig. 7f-f.”,j-j”). While the midline-crossing of spinal cord cells has been proposed to confer a morphogenetic advantage for zebrafish spinal cord development^18,21,25^, the regulation of this morphogenetic behavior is poorly understood. In particular, it is not clear whether this behavior is linked to neurogenic Notch signaling. We decided to address this issue in *mib1^ta52b^* mutants.

To visualize the midline-crossing of neural tube cells, one blastomere of two cell stage embryos was injected with GAP43-GFP RNA (Fig. 8a-h, Supplementary Fig.6). By the end of gastrulation, embryos in which the progeny of the injected blastomere occupied only the left or the right half of the embryo were selected for further analysis. Midline-crossing of neural tube cells is most prevalent from 14 to 18 hpf^17,20^. Accordingly, extensive crossing of GAP43-GFP positive cells to the contralateral side of the neural tube is observed by the 14 somites stage (i.e. 16 hpf) in wild-type siblings (Supplementary Fig.6a,b). In contrast, midline crossing is severely reduced in *mib1^ta52b^* mutants (Supplementary Fig.6c,d). Additional experiments confirmed that midline crossing is still reduced at later developmental stages (Fig. 8c,d, Supplementary Fig. 6e-h), establishing that this phenotype is not due to a developmental delay in mutant embryos.

**Figure 8:**
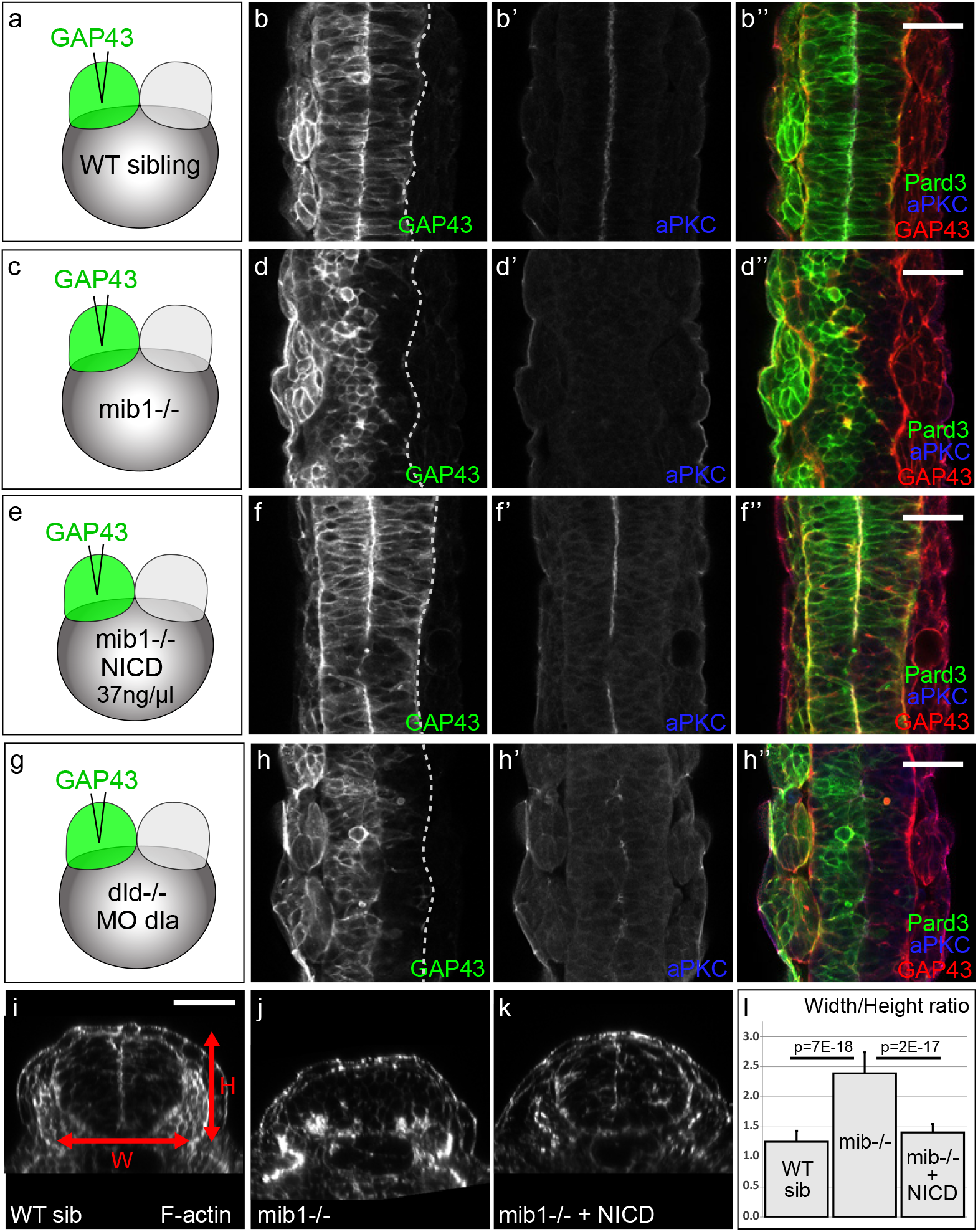
Notch signaling governs morphogenetic cell movements in the zebrafish spinal cord. (a-h) To label one half of the neural tube, RNA encoding a fluorescent membrane label (GAP43-GFP) was injected into one blastomere of two cell stage embryos (see Methods). (a,b) In WT siblings, cells from one half of the neural tube cross the organ midline to integrate the contra-lateral half (b,b”, n=9). (c,d) In 12/13 *mib1* mutants, neural tube cells fail to display this morphogenetic behavior. (e,f) RNA injection of constitutively activated Notch (NICD) restores apico-basal polarity (note apical aPKC enrichment at the neural tube midline in f’ compared to d’) and midline-crossing cell movements (f,f”) in 15/16 *mib1* mutants. (g,h) Neural tube cells display reduced midline crossing in 30/45 *dld* ; *dla* compound mutant/morphants. (i-l) Transversal sections of the anterior spinal cord were used to measure the width (W in i) and Height (H in i) of the neural tube. *mib1* mutants (i, n=14) present an increased W/H ratio compared to WT siblings (j, n=14). NICD RNA injection allows to restore neural tube proportions (k, n=25). (l) Mean W/H ratios for different conditions, error bars indicate standard deviation. Additional numerical informations are provided in the methods section. b,d,f,h are dorsal view of the anterior spinal cord at the 16 somites stage, anterior up. i-k are transversal sections of the anterior spinal cord at 30 somites, a stage when midline crossing is completed in WT embryos. Scalebars: 40 μm.

NICD injection allowed to restore midline crossing in *mib1^ta52b^* mutants (Fig. 8e,f), establishing thereby that this morphogenetic defect is due to a loss of Notch signaling and not to additional, Notch-independent, functions of Mib1 in the regulation of cell migration^33^. Accordingly, midline crossing is also reduced in *dld; dla* deficient embryos (Fig. 8g,h).

While the loss of Notch signaling activity impairs both the apico-basal polarization and the midline-crossing of neural tube cells, our experiments suggest that these two phenotypes are not strictly interdependent. Indeed, the injection of a low dose of NICD does not restore neural tube apico-basal polarity, but is sufficient to rescue midline crossing (Supplementary Fig. 6i,j).

The midline crossing of neural tube cells results in the intercalation of cells originating from the two sides of the neural tube, a behavior remindful of convergence extension movements^20,24^. We therefore wondered whether the shape of the spinal cord is altered in *mib1^ta52b^* mutants? In accordance with this hypothesis, transversal sections of the anterior spinal cord reveal an increased Width-to-Height ratio in *mib1^ta52b^* mutants (Fig. 8i,j,l). NICD injection restores neural tube proportions (Fig. 8k,l), conforming thereby that Notch signaling is required for the execution of morphogenetic movements that direct the proper shaping of the spinal cord.

## DISCUSSION

The aim of the present study was to investigate how cell fate specification and morphogenesis are linked during the development of the zebrafish nervous system. In higher vertebrates, a dual relationship exists between the polarized epithelial organization of the neural plate and neurogenic Notch signaling. Notch activation promotes the maintenance of polarized radial glia cells^10^ while the epithelial architecture of the neural primordium is itself required for Notch signaling^16^. In contrast, the early zebrafish neural plate does not display hallmarks of apico-basal polarity and the neural tube acquires a neuroepithelial tissue organization only when neurogenesis is well under way^17,18^. Our work shows that in spite of these apparent differences, Notch signaling is required for zebrafish spinal cord morphogenesis.

Previous studies have implicated noncanonical, transcription-independent Notch signaling in the late maintenance of apico-basal polarity in the ventral neural tube^42^. In contrast, we show here that the E3-Ubiquitin ligase Mib1, a critical regulator of Delta internalization and Notch activation^6^, is required to initiate the epithelialization of the neural primordium, allowing thereby the formation of a neural tube whose neuroepithelial organization is similar to the one of tetrapods (Fig. 1, Fig. 4).

Beyond the control of Delta ligand endocytosis, Mib1 has been shown to inhibit Epb41l5, a protein that facilitates the disassembly of apical junctional complexes^32^. Likewise Neuralized, which promotes Delta internalization in *Drosophila*, exerts a Notch-independent activity in epithelial morphogenesis^50^. Our observations show however that in the zebrafish spinal cord not only Mib1 itself, but the complete canonical Notch pathway, including its transcriptional mediators RBPJ/Su(H) are required for neuroepihtelial morphogenesis (Fig. 2). While polarized neuroepithelial cells are already present in higher vertebrates before the beginning of neurogenesis^13^, our findings show that in the zebrafish spinal cord neurogenic Delta/Notch signaling is itself required to promote neuroepithelial tissue organization.

Notch inactivation in the dorso-medial spinal cord causes excessive neuronal differentiation and the loss of neuroepithelial gene expression and polarity (Fig.5, Supplementary Fig.4,). It remains to be established whether Notch actively induces neuroepithelial properties or if, alternatively, these characteristics emerge by default as soon as Notch inhibits neuronal differentiation. As available tools do not allow manipulating Notch and neurogenic differentiation independently of each other, it is currently not possible to address this question.

Various mechanisms have been shown to govern the establishment of apico-basal polarity in different model systems^47^. Accordingly, Notch signaling is required for the morphogenesis of the dorso-medial spinal cord, but other regions of the nervous system can acquire neuroepithelial characteristics independently of Notch signaling (Fig.3, Fig.6, Supplementary Fig.5). These differences are likely due to the fact that the importance of Notch for developmental cell fate decisions varies according to the biological context. In contrast to more dorsal spinal cord cells, floor plate cells give rise to essentially glial derivatives^51^. Accordingly, *mib1* mutant floor plate cells do not undergo neuronal differentiation and retain their apico-basal polarity and expression of the radial glia marker *gfap*. At the level of the MHB, neurogenic differentiation is inhibited by the hairy-related transcription factor *her5* which acts independently of Notch signaling^52^, a situation that is different from the spinal cord where neurogenesis is regulated be the Notch-responsive *her4* gene^53^.

Spinal cord development requires not only neurogenesis, but also the execution of specific morphogenetic movements^18,19,21^. Zebrafish cells from one side of the neural tube invade the contralateral organ half by undergoing midline-crossing C-divisions^17,24,25^. We show that Notch signaling, which is critical for the regulation of neurogenesis, is also required for the midline-crossing behavior of neural tube cells (Fig.8). Low levels of Notch signaling restore midline crossing but not apico-basal polarization of neural tube cells suggesting however that these two aspects of cellular behavior can be, at least partially, uncoupled (Supplementary Fig.6i).

Manipulations of PCP have been shown to impair the midline-crossing behavior of neural tube cells^20,24^. Due to the overall impact of PCP on embryonic convergence-extension movements, these experiments have however not allowed to evaluate the actual impact of this morphogenetic behavior on the shaping of the neural tube. We show that in *mib1^ta52b^* mutants, which do not display general convergence extension phenotypes, the cells of the neural primordium fail to display midline crossing and give rise to a misproportioned spinal cord (Fig.8i-l).

In conclusion, our findings show that, in addition to regulating neuronal cell fate specification, Notch signaling is essential to control neuroepithelial tissue organization and morphogenetic movements during the development of the zebrafish spinal cord. In the future, it will be interesting to further characterize the molecular mechanisms through which the Notch pathway governs the morphogenesis of the nervous system.

## METHODS

### Zebrafish strains and genotyping

Zebrafish strains were maintained under standard conditions and staged as previously described^54^. Zebrafish embryos were grown in 0.3x Danieau medium.

To genetically inactivate *mindbomb1*, we used the *mib1^ta52b^* allele^6^. Depending on the experiment, three different criteria were used either separately or in combination to identify *mib1^ta52b^* homozygous animals: 1, DeltaD immunostaining (mutant embryos can be identified by upregulated DlD signal at the cell membrane, Fig. 1c”); 2, embryo morphology; and 3, molecular genotyping. For the latter, a 4-primer-PCR was used to identify *mib1^ta52b^* and WT alleles in a single PCR reaction. The following primers were used: 5’-ACAGTAACTAAGGAGGGC-3’ (generic forward primer), 5’-AGATCGGGCACTCGCTCA-3’ (specific reverse primer for the WT allele), 5’-TCAGCTGTGTGGAGACCGCAG-3’ (specific forward primer for the *mib1^ta52b^* allele), and 5’-CTTCACCATGCTCTACAC-3’ (generic reverse primer). PCR amplification was carried out using GoTaq polymerase (Promega) at 1.5 mM MgCl2 using the following cycling parameters: 2 min 95°C - 10 cycles [30 sec 95°C – 30 sec 65 to 55°C – 60 sec 72°C] – 25 cycles [30 sec 95°C – 30 sec 55°C – 60 sec 72°C] – 5 min 72°C. WT and *Mib1^ta52b^* mutant alleles respectively yield 303 bp and 402 bp amplification fragments. As some zebrafish strains present polymorphic *mib1* WT alleles, it is important to validate the applicability of this protocol before using it in a given genetic background.

To inactivate *deltad* we used *dld/aei^AR33^*^55^ mutant embryos obtained through incrossing of homozygous mutant adult fish. To visualize Notch signaling activity, we used the *tp1bglob:eGFP* transgenic line^44^.

### mRNA and morpholino injections

Microinjections into dechorionated embryos were carried out using a pressure microinjector (Eppendorf FemtoJet). Capped mRNAs were synthesized using the SP6 mMessage mMachine kit (Ambion) and poly-adenylated using a polyA tailing kit (Ambion). RNA and morpholinos were injected together with 0.2% Phenol Red.

RNA microinjection was performed using the following constructs and concentrations: Mindbomb1-pCS2+ (125 ng/μl)^34^; Pard3-GFP-pCS2+ (50 ng/μl)^29^; DN-Su(H)-pCS2+ (600 ng/μl)^39^; CA-Su(H)-pCS2+ (40ng/μl)^39^; Myc-Notch-Intra-pCS2+ (25-37.5 ng/μl)^38^; Gap43-GFP-pCS2+ (20 ng/μl) and GAP43-RFP-pCS2+ (30 ng/μl).

Morpholino oligonucleotides were injected at the indicated concentrations to knock down the following genes: mindbomb1: 5’ -GCAGCCT CACCT GTAGGCGCACTGT-3’ (1000μM)^6^; deltaA: 5’- CTTCTCTTTTCGCCGACTGATTCAT-3’ (250 μM)^35^; RBPJa: 5’-GCGCCATCTTCACCAACTCTCTCTA-3’ (50 μM) and RBPJb: 5’-GCGCCATCTTCCACAAACTCTCACC-3’ (50 μM). To ensure that the phenotypes of *dla/dld^AR33^* morphant/mutants and RBPJa&b double morphants were not due to non-specific p53-mediated responses, we performed these experiments in the presence of a validated p53 Morpholino (5’- GCGCCATTGCTTTGCAAGAATTG-5’, 333μM)^56^.

### Gamma-secretase inhibitor treatment

At mid-gastrulation zebrafish embryos were transferred to 0.3x Danieau medium containing 50 μM LY411575^37^ (Sigma) or 100 μM DAPT^36^ (Sigma) dissolved in DMSO. Embryos were raised till the 30 somites stage before being processed for antibody staining. Control embryos were mock-treated with DMSO alone.

### Whole mount *in situ* hybridization

*In situ* hybridization was performed according to Thisse et al.^57^. DIG-labeled antisense RNA probes were transcribed from PCR products carrying the T7-promoter sequence (5’-TAATACGACTCACTATAGGG-3’) on the reverse primer. PCR amplicons for the different genes were flanked by the following sequences: *sox19a:* forward: 5’-CGATGTCGGGTGAAGATG-3’, reverse: 5’- CTGTCAAGGTTGTCAAGTCAC-3’*gfap:*forward: 5’-TAAAGAGTCCACTACGGAGAGG-3’, reverse: 5’-GGCACCACAATGAAGTAATGTCC-3’, *crumbs1:* forward: 5’-TGTACCACCAGCCCATGTCATA-3’, reverse: 5’-cctcatcacagttttgacccac-3’; *crumbs2a:* forward: 5’-TGAGAGTGCCCCCTGCCTTAAT-3’, reverse: 5’-acagtcacagcggtagc-3’; *pard6γb:* forward: 5’- GACTACAGCAACTTTGGCACCAGCACTCT-3’, reverse: 5’-gtgatgactgtgccatcctcctc-3’.

### Immunocytochemistry

Embryos were fixed in 4% paraformaldehyde in PEM (PIPES 80 mM, EGTA 5 mM, MgCl2 1 mM) for 1.5 hours at room temperature or overnight at 4°C, before being permeabilized with 0.2% TritonX-100 in PEM-PFA for 30 minutes at room temperature. Subsequent washes and antibody incubations were performed in PEM+0,2% TritonX-100. Primary antibodies used were: Mouse@DeltaD^6^ (1:500, Abcam ab73331); Mouse@DeltaA^58^ (1:250, ZIRC 18D2);Rabbit@aPKC^28^ (1:250, Santa Cruz sc-216); Mouse@ZO1^59^ (1:500, Invitrogen 1A12); Mouse@HuC/D^60^ (1:500, Invitrogen 16A11); Rabbit@γ-Tubulin (1:250, Sigma T5192).

### Cell transplantations

For cell transplantation embryos were maintained in 1x Danieau medium + 5% penicillin-streptomycin. Donor embryos were labelled by injection of RNA encoding Pard3-GFP at the one-cell stage. Cell transplantations were carried out at late blastula / early gastrula stages. In each experiment, 20-30 cells were aspirated from the donor embryo using a manual microinjector (Sutter Instruments) and transplanted into the host embryo. Transplanted embryos were grown till the 30 somites stage in agarose-coated petri dishes with 0.3x Danieau and 5% penicillin-streptomycin before being fixed and processed for antibody staining.

### Analysis of neural tube cell midline-crossing behaviour

To label one half of the neural tube, GAP43-GFP RNA was injected into one blastomere of 2-cell stage zebrafish embryos. The embryos were then grown till the bud stage, at which time point the localisation of the fluorescent cells was analysed using a fluorescent microscope (Leica M205 FA). In a typical experiment, about 50% of the embryos displayed a unilateral localisation of GFP-positive cells and were kept to be grown till the desired stage before being fixed and processed for antibody staining. In contrast to the cells of the neural tube, somitic precursors do not cross the embryonic midline in the course of development. Consequently, successful half-injection results in a unilateral labelling of the somites that becomes visible at confocal analysis.

To separately label the two opposite sides of the neural tube, 50 2-cell stage embryos were initially injected into one blastomere with RNA encoding the first fluorescent membrane label (e.g. GAP43-GFP). The presence of Phenol red in the injection mix allows identifying the injected blastomere for several minutes after injection. A second injection needle was then used to inject the second RNA (e.g. GAP43-RFP) into the other blastomere. Embryos were grown till the bud stage and screened for efficient double half injection as described above.

### Microscopy and image analysis

For confocal imaging, embryos were mounted in 0.75% low melting agarose (Sigma) in glass bottom dishes (MatTek corporation). Embryos were imaged on Spinning disk (Andor) or Laser scanning confocal microscopes (Zeiss LSM510, 710, 780 and 880) using 40x Water or 60x Oil immersion objectives. *In situ* gene expression patterns were documented on a Leica M205FA-Fluocombi stereomicroscope. Image analysis was performed using ImageJ (http://rbs.info.nih.gov/ii/) or Zeiss ZEN software.

For the quantification of the temporal progression of apico-basal polarity in the neural tube (Fig. 4M), we acquired confocal stacks spanning the entire dorso-ventral extent of the neural tube. The percentage of neural tube polarity was then calculated for each embryo as the number of confocal slices displaying polarized aPKC enrichment, divided by the total number of slices of the neural tube stack. Error bars in Fig. 4M represent standard deviations. Examples of individual confocal slices at different dorso-ventral locations are shown in Fig. 4A-L. Due to the thickness of the neuro-epithelial cells that constitute the dorsal and ventral walls of the neural tube, the neural tube lumen (and therefore the measurable extent of polarized aPKC) never reaches the full dorso-ventral extent of the neural tube (which would correspond to 100% in Fig. 4M).

For the analysis of the spinal cord Width-to-Height (W/H) ratio displayed in Fig. 8i-l, embryos were stained with fluorescent Phalloidin and DAPI and mounted in glass bottom dishes with the dorsal surface of the embryo facing the coverslip. Embryos were imaged using a Zeiss LSM880 confocal microscope. A line scan was performed at the border between the 2^nd^ somite and the 3^rd^ somite to obtain a transversal section of the spinal cord. Zeiss ZEN imaging software was used to measure the width and the height of the spinal cord before calculating the width-to-height ratio in Excel. The W/H ratios for the indiviual display items in Fig.8 are WT sib (i) 1.249, mib1-/- (j) 2.285, mib1-/- + NICD (k) 1.409. The Mean W/H ratios ± SD for the graph displayed in Fig. 8l are WT 1.253 ± 0.183, mib1-/- 2.391 ± 0.347, mib1-/- + NICD 1.407 ± 0.140.

### Statistical analysis

For the analysis of spinal cord Width-to-Height (W/H) ratio reported in Fig. 8i-l, statistical analysis was carried out using the R/RStudio packages. The normal distribution of experimental data was verified using the Shapiro-Wilk test. Pairwise comparisons of the data sets displayed in Fig. 8l were performed using a pairwise t-test with Bonferroni correction for multiple comparisons.

### Use of research animals

Animal experiments were performed in the iBV Zebrafish facility (experimentation authorization #B-06-088-17) in accordance with the guidelines of the ethics committee Ciepal Azur and the iBV animal welfare committee.

### Data availability

The datasets generated during analysed during the current study are available from the corresponding author on reasonable request.

## ACKNOWLEDGEMENTS

This study was supported by a CNRS/INSERM ATIP/Avenir 2010 grant and an HFSP Career Development Award (00036/2010) to M.F. P.S. benefited from an FRM 4^th^ year PhD fellowship (FDT20140930987). L.X. was supported by an ARC postdoctoral fellowship (PDF20121206203). V.S. is supported by the LABEX SIGNALIFE PhD program (ANR-11-LABX-0028-01). Confocal microscopy was performed with the help of the iBV PRISM imaging platform. We thank L. Bally-Cuif, C. Kinter, E.Ober, Y.J. Jiang, J. Malicki, S.Sokol, A.Oates and D. Stainier for the sharing of reagents. We are grateful to S. Polès, M.A. Derieppe, R. Rebillard and F. Paput for excellent technical assistance.

## AUTHOR CONTRIBUTIONS

P.S. performed the experiments reported in Fig.1a-g, Fig.2a-f,m,n, Fig.4, Fig.5i-t, Fig.6, Fig.7a,b and Fig.S2a-h. V.S. performed the experiments of Fig. 1h-k, Fig.2i-l,o,p, Fig.7c-j, Fig.8 Fig.S2i-l and Fig.S6. L.X. performed the experiment displayed in Fig. 2g,h. M.F. provided the data for Fig.3, Fig.5a-h, Fig.S1, Fig.S3, Fig.S4a-d and Fig.S5. M.F. designed and supervised the study and wrote the manuscript. All authors analyzed the data and contributed to the final version of the manuscript.

## ADDITIONAL INFORMATION

### Supplementary Information

Supplementary Information includes six figures.

---

**Supplementary Figure 1.**
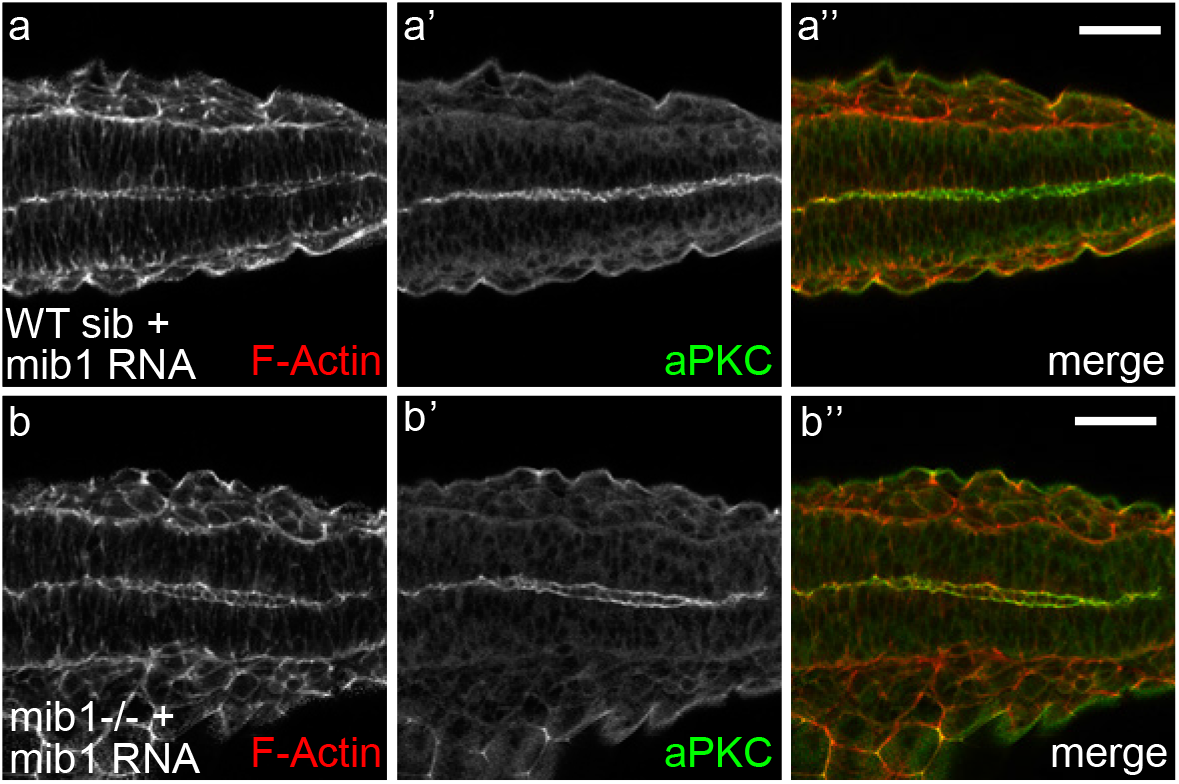
Mindbomb is important for zebrafish spinal cord morphogenesis. (a,b) Injection of RNA encoding wild-type Mib protein restores neuroepithelial morphology (visualized using F-Actin staining) and polarized accumulation of the apical marker aPKC in *mib1* mutants (5/7 embryos full rescue, 2/7 partial rescue). Dorsal views of the anterior spinal cord at the 18 somites stage, anterior left. Scalebars: 50 μm.

**Supplementary Figure 2.**
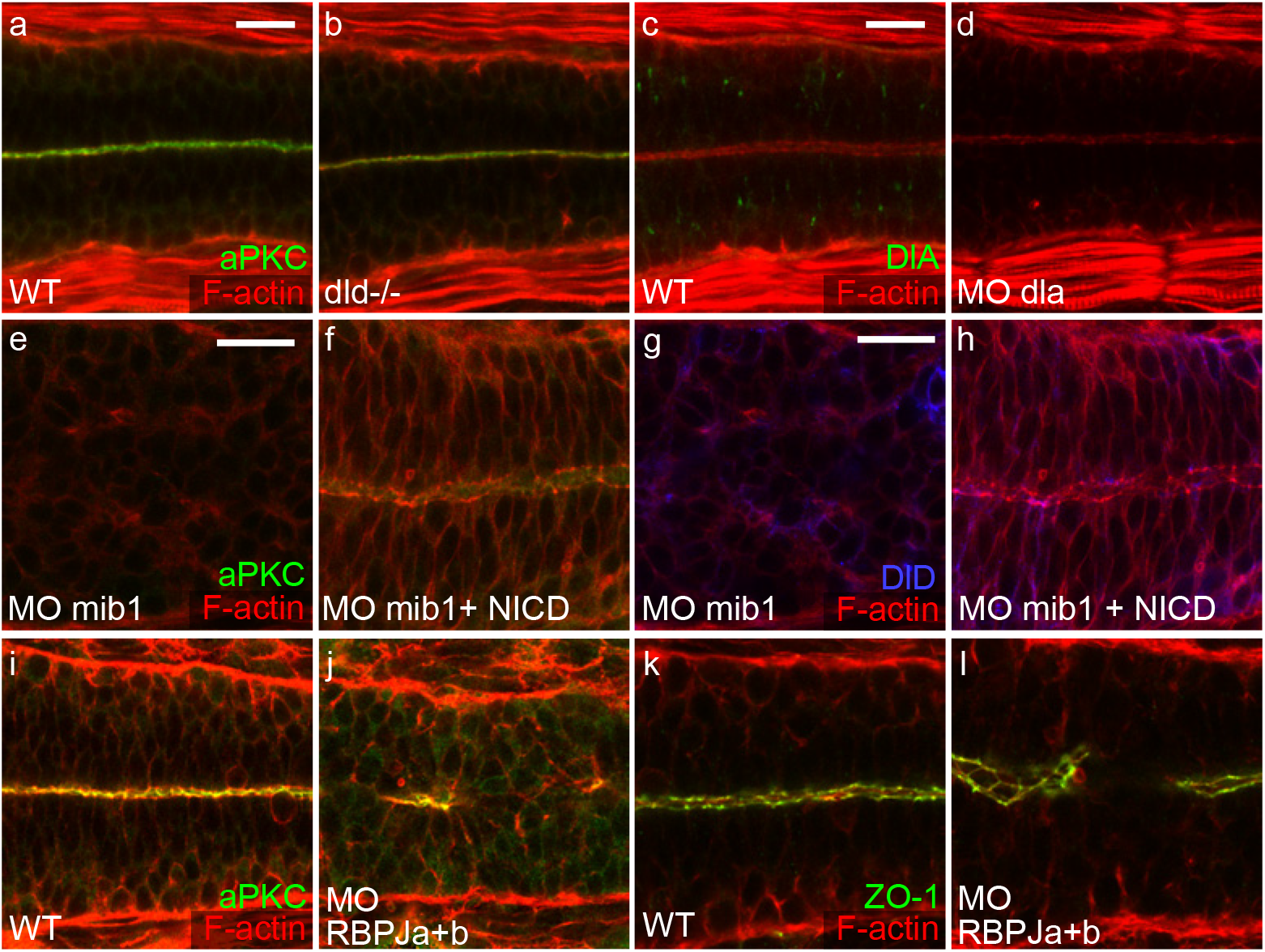
Notch pathway components are required for the apico-basal polarity of the zebrafish spinal cord. (**a,b**) Apical aPKC staining and cortical F-Actin indicate normal apico-basal polarity and neuroepithelial morphology in *d1d^AR33^* mutants (n=8/8). (**c,d**) dlA morpholino injection does not alter neuroepithelial morphology but abolishes DlA immuno-reactivity (n=4/4). (**e-h**) RNA injection of a constitutively activated form of Notch (NICD) restores neuroepithelial morphology and apical aPKC localisation (**f**) but not DeltaD endocytosis (**h**) in 14/17 embryos. (**i-l**) Embryos injected with morpholinos against RBPJa & b display a partial loss of apico-basal polarity as visualized by a partial disruption of apical aPKC (**j**, n=23/29) and ZO-1 (**l**, n=16/16). (**a-d**) 26 somites stage, (**e-l**) 30 somites stage. Phalloidin staining of F-Actin is used to highlight cell outlines. Scalebars: 20 μm.

**Supplementary Figure 3.**
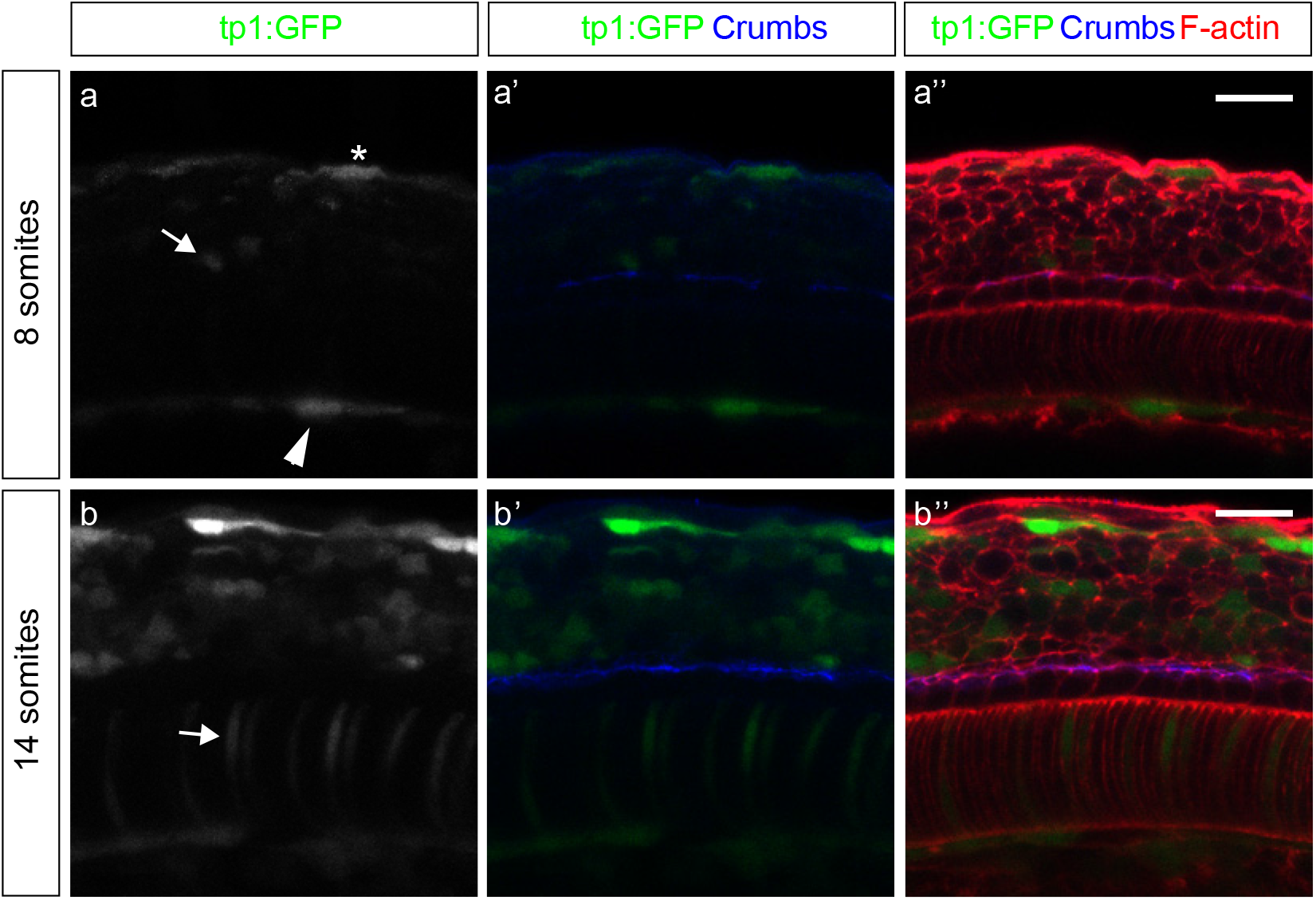
Early floor plate morphogenesis occurs in the absence of detectable Notch reporter activity. (**a,b**) Lateral views of floor plate cells at the level of the anterior spinal cord, anterior to the left, dorsal up. (**a**) At the 8 somites stage, the activity of the transgenic Notch reporter line *tp1:bglob-GFP* (tp1:GFP) is detected in isolated neural tube cells (arrow), hypocord cells (arrowhead) and epidermal cells (star) (n=13). (**b**) At 14 somites, numerous GFP-positive cells are found in the neural tube and additional reporter expression is detected in notocord cells (arrow) (n=13). Floor plate cells, which can be identified through their cuboidal morphology (F-Actin in **a”,b”**) do not display Notch reporter activity. 8 and 14 somites stage embryos were imaged using the same confocal settings. For display purposes, contrast enhancement was then used to improve the visibility of the weak 8 somites stage *tp1:GFP* and Crumbs signals in **a-a”**. Scalebars: 20 μm.

**Supplementary Figure 4.**
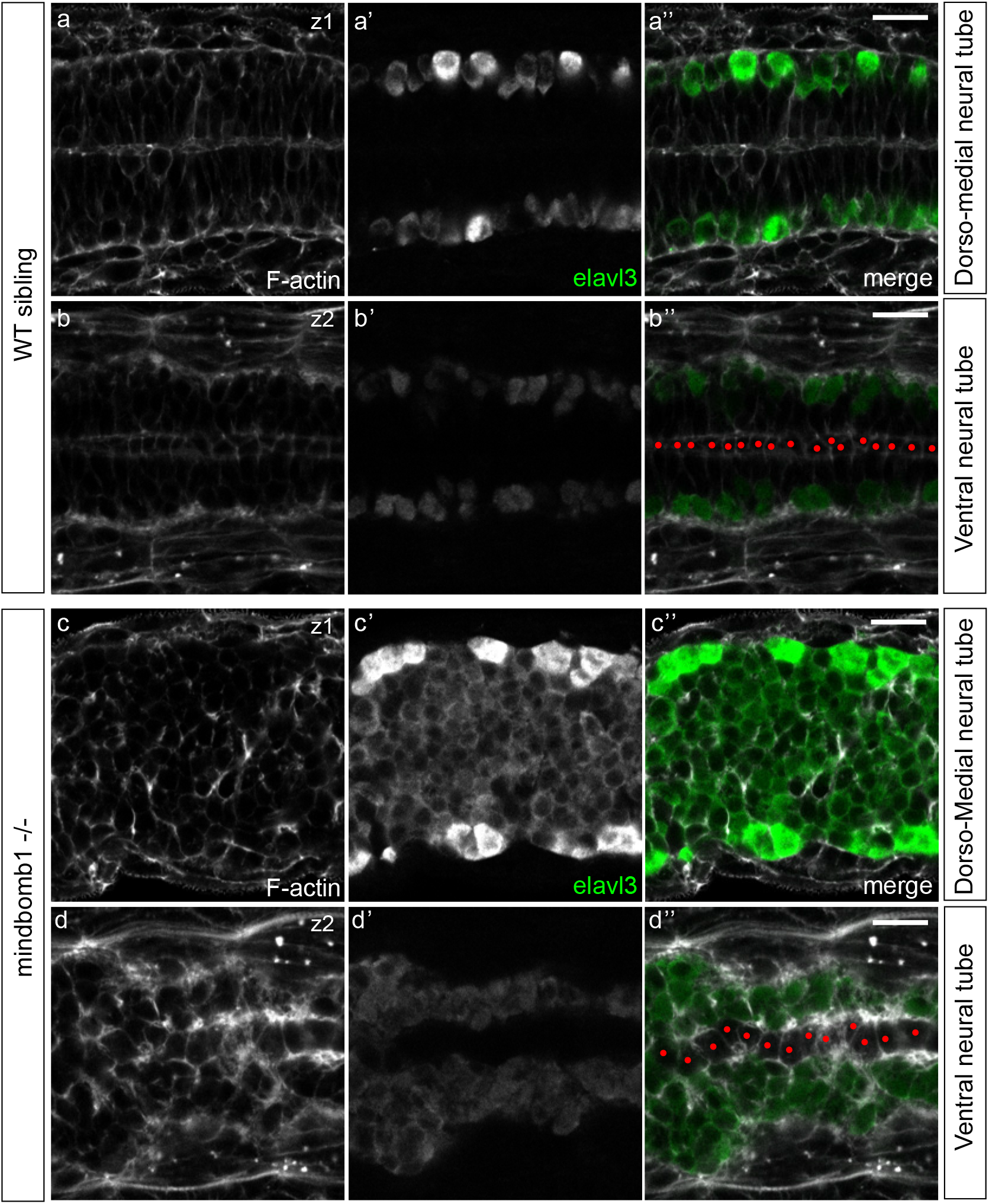
Excessive neuronal differentiation causes neural tube morphogenesis defects in *mindbomb1* mutants. (**a-d**) In 22 somites stage WT siblings, the neuronal differentiation marker *elavl3a* is expressed in some cells of the dorso-medial (**a’,a”**) or ventral (**b,b”**) spinal cord. In *mib1* mutants, all dorso-medial neural tube cells differentiate as neurons (**c’,c”**), causing a loss apico-basally polarized precursor cells and a disruption of neuro-epithelial morphology (n=4/4). In the ventral neural tube, floor plate cells can be identified by their cuboidal morphology (red dots in **b”,d”**). In *mib1* mutants, floor plate cells are the only ones not undergoing neuronal differentiation (**d’,d”**, n=4/4). a,b and c,d are views of the same embryo at different z levels. All images are dorsal views of the anterior spinal cord, anterior left. Scalebars: 20 μm.

**Supplementary Figure 5.**
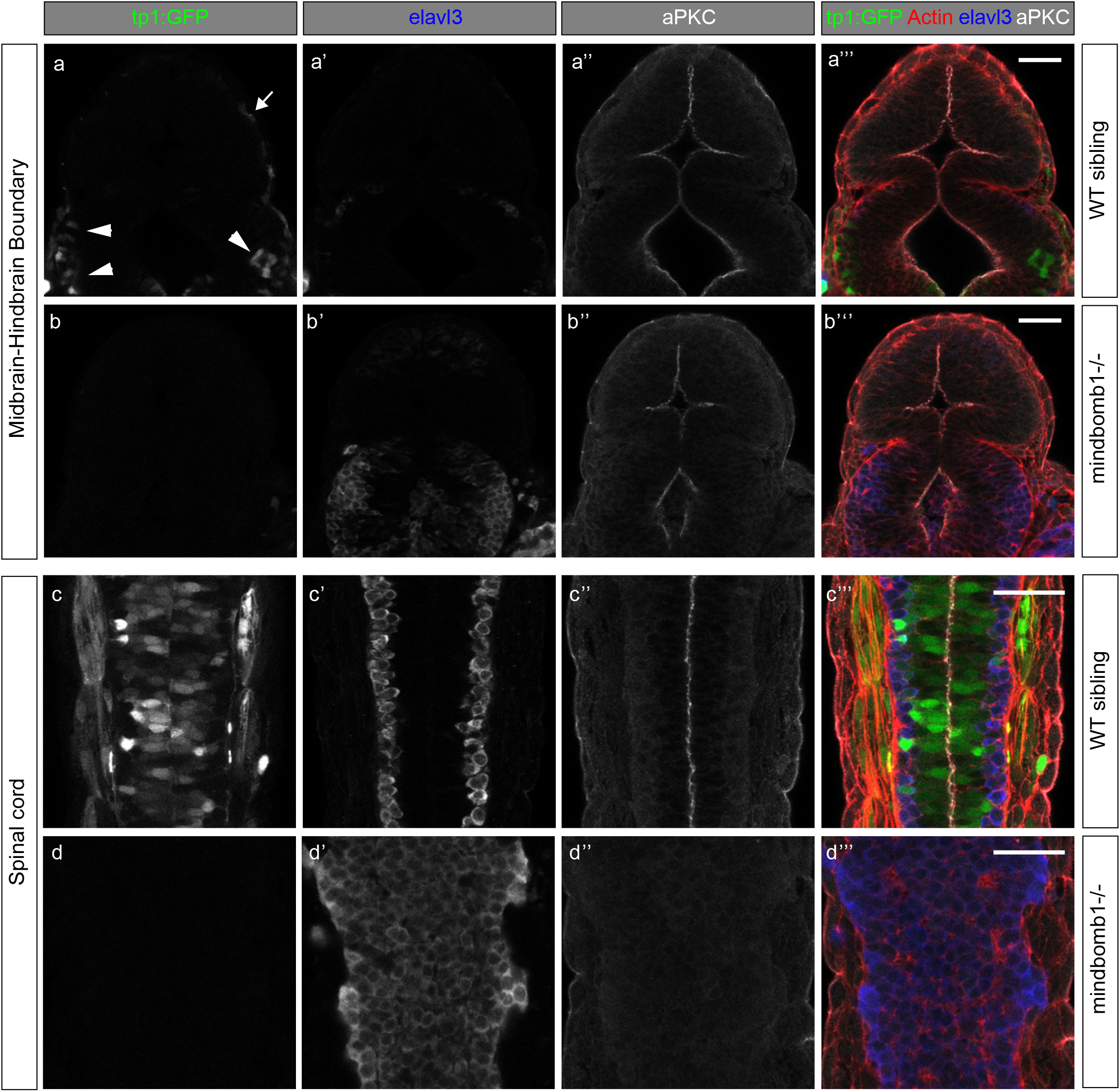
Tissue-specific regulation of neurogenesis and neuro-epithelial tissue organization. (**a-d**) Dorsal views of the Midbrain-Hindbrain Boundary (**a,b**) and anterior spinal cord (c,d) at the 30 somites stage, anterior up. In WT siblings, the *tp1bGlob:GFP (tp1:GFP)* Notch reporter transgene indicates active signaling in a small number of cells in the anterior hindbrain (arrowheads in a). More anteriorly, Notch activity is detected only in epidermal cells (arrow in a) but not in the midbrain itself. *mib1* mutants present excessive neurogenesis (**b’**) and a partial disruption of apical aPKC localisation (**b”**) at the level of the anterior hinbrain (n=9). Only few elavl3-positive neurons are detected in the midbrain, which maintains a proper neuroepithelial organization (**b”,b”’**). (**c,d**) At the level of the anterior spinal cord, WT sibling embryos display widespread Notch reporter activity (**c**), basaly localized elavl3-positive neurons (**c’**) and polarized aPKC enrichment at the apical neural tube midline (**c”**). *mibl* mutants present a loss of Notch reporter expression (**d**), excessive neuronal differentiation (**d’**) and a lack of polarized aPKC localization (**d”**) (n=9). a,c and b,d respectively represent different regions of the same WT and *mib1* mutant embryos. Scalebars: 40 μm.

**Supplementary Figure 6.**
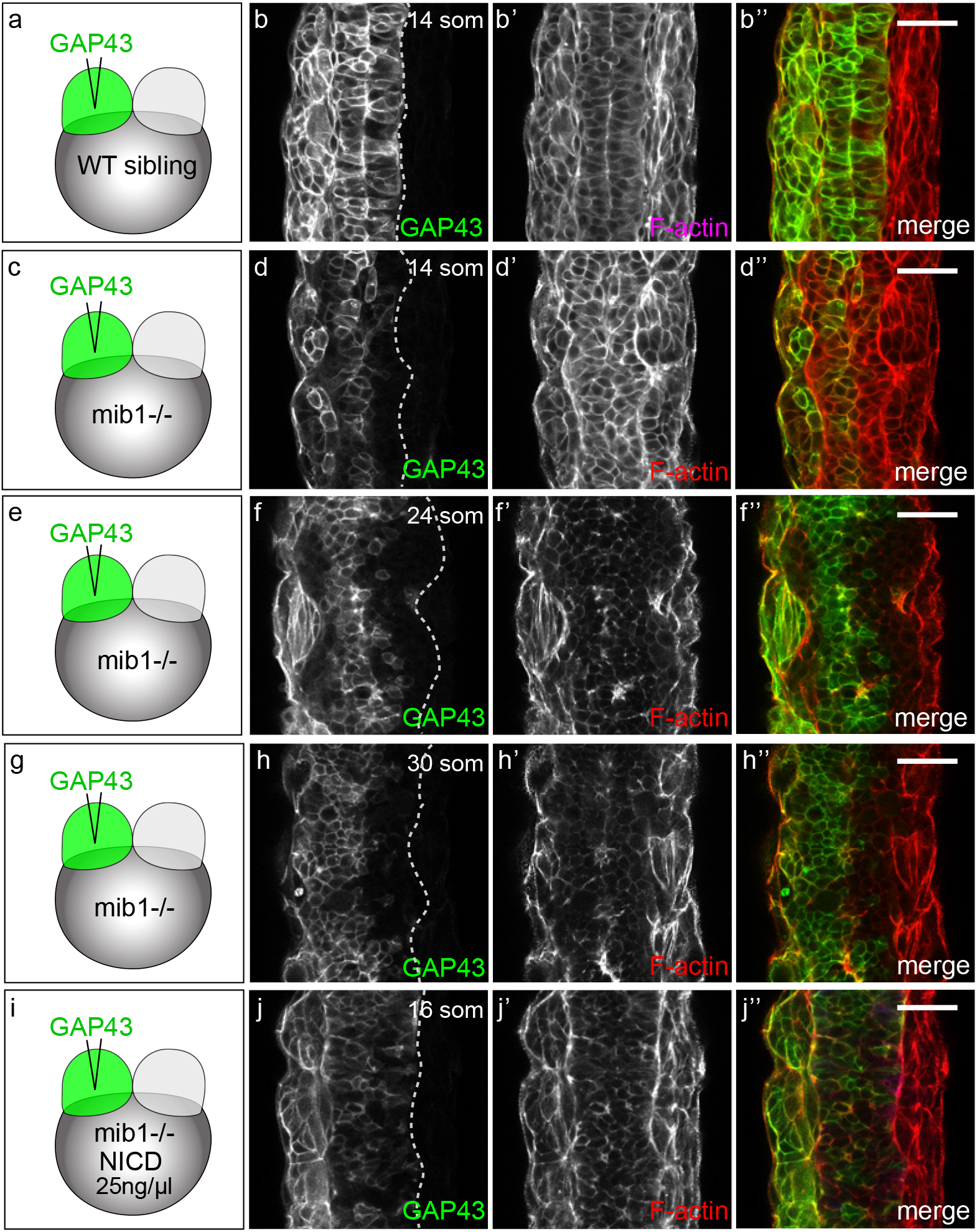
*Mindbomb1* loss of function impairs the midline-crossing behavior of neural tube cells. (**a-j**) One half of the neural tube was labelled by injecting RNA encoding a green membrane label (GAP43-GFP) into one blastomere of 2-cell stage embryos (see Methods). (**a,b**) In 5/5 WT sibling embryos at the 14 somites stage, cells originating from one side of the neural tube have crossed the neural tube midline to integrate the contra-lateral organ half. (**c,d**) In contrast, cells fail to cross to the contra-lateral side in 8/8 *mib1* mutants. (**e-h**) This inhibition of midline-crossing persists if *mib1* mutants are analysed a the 24 somites (**e,f**, n=11/13) or 30 somites (**g,h**, n=11/13) stages. (**i,j**) Injection of a low dose of constitutively activated Notch (NICD) that does not restore neuroepithelial tissue architecture (note the lack of apical F-actin accumulation at the neural tube midline in **j’**) is sufficient to restore midline-crossing in 5/7 *mib1* mutants. All images are dorsal views of the anterior spinal cord, anterior up. Scalebars: 40 μm.

